# Loss of H3K9 tri-methylation alters chromosome compaction and transcription factor retention during mitosis

**DOI:** 10.1101/2022.02.01.478684

**Authors:** Dounia Djeghloul, Andrew Dimond, Holger Kramer, Karen Brown, Bhavik Patel, Yi-Fang Wang, Matthias E. Futschik, Chad Whilding, Alex Montoya, Nicolas Veland, Sherry Cheriyamkunnel, Thomas Montavon, Thomas Jenuwein, Matthias Merkenschlager, Amanda G. Fisher

## Abstract

Recent studies have shown that repressive chromatin machinery, including DNA methyltransferases (DNMTs) and Polycomb Repressor Complexes (PRCs), bind to chromosomes throughout mitosis and their depletion results in increased chromosome size. Here we show that enzymes that catalyse H3K9 methylation, such as Suv39h1, Suv39h2, G9a and Glp, are also retained on mitotic chromosomes. Surprisingly however, mutants lacking H3K9me3 have unusually small and compact mitotic chromosomes that are associated with increased H3S10ph and H3K27me3 levels. Chromosome size and centromere compaction in these mutants were rescued by providing exogenous Suv39h1, or inhibiting Ezh2 activity. Quantitative proteomic comparisons of native mitotic chromosomes isolated from wildtype versus Suv39h1/Suv39h2 double-null ESCs revealed that H3K9me3 was essential for the efficient retention of bookmarking factors such as Esrrb. These results highlight an unexpected role for repressive heterochromatin domains in preserving transcription factor binding through mitosis, and underscore the importance of H3K9me3 for sustaining chromosome architecture and epigenetic memory during cell division.

## Introduction

Heterochromatin and euchromatin are defined cytologically as condensed and decondensed regions of the genome, respectively^1, 2^. Constitutive heterochromatin, although gene-poor, contains non-coding DNA repeat elements that are often abundantly transcribed^3, 4^. During interphase, heterochromatin-containing domains within different chromosomes cluster together forming structures termed chromocenters^5–8^. In mouse cells these dynamic structures are characteristically marked by a high density of histone H3K9me3 and histone H4K20me3, as well as the H3K9-associated heterochromatin protein 1 (HP1α or Cbx5)^1, 7^. As chromosomes condense and enter mitosis, primary constrictions first become evident within the heterochromatin domains that correspond to centromeric microsatellite (MiSat) arrays, where the mitotic spindles will eventually bind. These centromeric regions are flanked by much larger domains of pericentric non-coding major satellite repeats (MaSat) that in the mouse account for approximately 3.6% of the genome^9–11^.

The Suv39h1 and Suv39h2 histone lysine methyltransferases are hallmark enzymes of mammalian heterochromatin that catalyse the trimethylation of histone H3 at lysine 9 (H3K9)^12–14^. They form part of a larger group of enzymes capable of modifying H3K9, which include G9a (Ehmt2), Glp (Ehmt1)^15, 16^, Setdb1 and Setdb2^17, 18^, that primarily mediate H3K9 mono- and di-methylation, as well as members of the Kdm1, Kdm3 and Kdm4 families of histone demethylases^19–23^. While heterochromatin regulation is recognised as being essential in preserving nuclear architecture, genome stability, DNA repair and for silencing transposon expression in early mouse development^24–27^, the underlying repetitiveness of satellite DNA means that factors binding to heterochromatin are often excluded from conventional analyses.

In this study we set out to examine the impact of repressive H3K9me3 on mitotic chromosome architecture and on the factors that bind chromosomes through mitosis. Several DNA-binding transcriptional regulators have been shown to remain bound to chromosomes during mitosis, including Foxa1, Esrrb, Sox2 and Gata1^28–33^, where they occupy a subset of the genomic sites that are present during interphase. Such demonstrations have raised the interesting possibility that ‘mitotic bookmarking’ by factors such as these could help to convey cellular identity to newly divided daughter cells. Recent studies have also shown that many components of repressive chromatin machinery, including those that characterise constitutive heterochromatin, are also retained at mitotic chromosomes during cell division^34–36^. While this has prompted speculation of an interplay between mitotic booking factors and repressive chromatin states^37^, there is currently a paucity of direct evidence to support this. Here we confirm that factors mediating H3K9 trimethylation are indeed retained on native mitotic chromosomes isolated from different cell types. H3K9me3 depletion results in altered chromosome architecture, compensatory changes in the level and distribution of repressive H3K27me3, and discrete and reversible changes in the retention of specific mitotic bookmarking factors.

## Results

### Reduced chromosome size and increased centromere compaction are features of *Suv39h dn* mitotic chromosomes

Prior studies had suggested that Suv39h1, Suv39h2 and HP1 proteins (Cbx5 or HP1α, Cbx1 or HP1β, and Cbx3 or HP1ɣ), as well as other repressive chromatin machinery, remain bound to chromosomes during mitosis^34–36, 38–40^. Suv39h1, Suv39h2 and HP1 enrichment at mitotic chromosomes was independently confirmed herein using established proteomic approaches^36^ in mouse ESCs and mouse embryonic fibroblasts (MEFs) (outlined in Figure S1a and depicted in S1b). To investigate the impact of H3K9me3 on mitotic chromosome architecture, native mitotic chromosomes were isolated directly from wild type (WT) ESCs (Figures 1a-c) or MEFs (Figures 1d-f) and compared with mitotic chromosomes isolated from cells lacking both Suv39h1 and Suv39h2 (Suv39h double null or *Suv39h dn*)^13, 26^. To enable this, dividing cell cultures of ESCs were arrested in metaphase using demecolcine, chromosomes released and stained with Hoechst 33258 and Chromomycin A3 and individual chromosomes isolated by flow cytometry as described previously^36^ (Figure S1a provides a schematic of the approach). Using purified preparations of chromosomes 19 and X as exemplars, mitotic chromosomes isolated from *Suv39h dn* ESCs^26^ were shown to be smaller than their WT counterparts, and a marked increase in compaction at centromeric domains was noted (Figures 1b and 1c).

**Figure 1.**
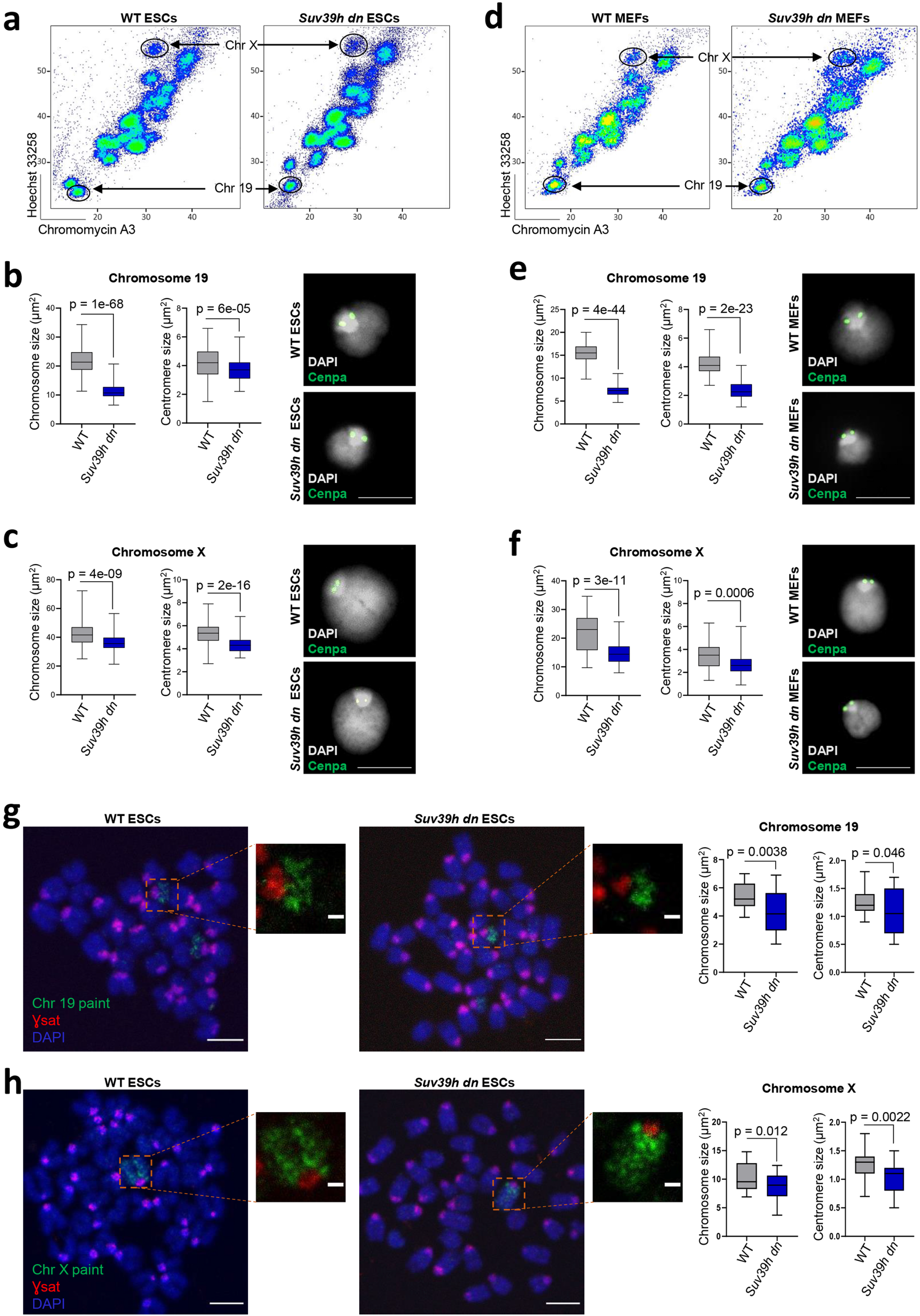
Suv39h1/Suv39h2 deficiency generates small and compact mitotic chromosomes. (a-f) Flow sorting and size measurements of chromosomes 19 and X from WT or *Suv39h dn* mouse ESCs and MEFs. Flow karyotype of mitotic chromosomes isolated from WT or *Suv39h dn* ESCs (a) and MEFs (d); gates used to isolate chromosomes 19 and X are indicated. Representative images (right) of mitotic chromosomes 19 (b,e) and X (c,f) from WT or *Suv39h dn* ESCs (b,c) and MEFs (e,f) are shown, where DAPI stain (grey) and Cenpa label (green) indicate the chromosome body and centromere respectively. Scale bars = 5 μm. Box plots (left of images) show area measurements of individual chromosomes and centromeres for WT and *Suv39h dn* cells. Minimum, lower quartile, median, upper quartile and maximum values are indicated. n = minimum 100 chromosomes analysed for each cell line over three independent experiments. P-values of statistically significant changes, measured by unpaired two tailed Student’s t-tests, are indicated. (g,h) Representative images of WT or *Suv39h dn* ESC metaphase spreads stained with chromosome 19 painting probe (green) (g), or with chromosome X painting probe (green) (h), in addition to gamma satellite probe (red) and DAPI (blue). Scale bars = 4 μm and 1 μm for the metaphase spread and zoom-in images respectively. Chromosome and centromere sizes of chromosomes 19 and X were calculated on metaphase spreads of WT and *Suv39h dn* ESCs. Box plots show chromosome and centromere area measurements for WT and *Suv39h dn* spreads. Minimum, lower quartile, median, upper quartile and maximum values are indicated. n = minimum 25 chromosomes analysed on metaphase spreads for each line over three independent experiments. P-values of statistically significant changes, measured by unpaired two tailed Student’s t-tests, are indicated.

To determine whether reduced size and enhanced condensation of mitotic chromosomes deficient in H3K9me3 was seen in other cell types, including differentiated cells, mitotic chromosomes from MEFs lacking Suv39h1 and Suv39h2^13^ were also examined (Figure 1d).

Consistent with results obtained in ESCs, mitotic chromosomes isolated from these fibroblasts were smaller than WT equivalents and showed increased compaction at centromeres (Figures 1e and 1f). To exclude that these changes were an artefact of the isolation procedure, we also examined conventional metaphase spreads where chromosome-specific probes and DNA FISH was used to label chromosomes 19 and X. These analyses confirmed that mitotic chromosomes from *Suv39h dn* ESCs were reproducibly smaller than their WT counterparts (Figures 1g and 1h). To examine whether deficits in H3K9 mono- and di-methylation affect chromosome compaction, we isolated mitotic chromosomes from a panel of MEFs that lacked Suv39h1 and Suv39h2 (CRISPR clone B1) or other H3K9 methyltransferases, specifically pairwise deletions of Setdb1 and Setdb2 (CRISPR clone A4 + 4-hyroxytamoxife or OHT), or G9a and Glp (CRISPR clone H7)^41^ (Figure S1c). Analysis of isolated native mitotic chromosomes from clones A4+OHT, H7, and B1 fibroblasts vs WT controls showed that reduced chromosome size was a unique feature of cells lacking H3K9me3^41^ (Figure S1d).

The compaction of mitotic chromosomes lacking H3K9me3 was unexpected as prior studies had shown that an absence of other repressive chromatin modifiers, such as DNA methylation or PRC2 activity, produced chromosomes that were larger and more decondensed than equivalent WT mitotic controls^36^. To investigate the possibility that additional heterochromatin modifications, such as increased H3K27me3, might compensate for deficits in H3K9me3, we examined the distribution of modified histones in *Suv39h dn* and WT ESCs. As anticipated, H3K9me3 decorated centromeric domains of normal mitotic chromosomes (Figures 2a and S2a green, WT top panel), but was absent from equivalent *Suv39h dn* chromosomes (Figures 2a and S2a, lower panel, quantified in the graph, left). Instead, we detected increased levels of H3K27me3 across centromeric domains of *Suv39h dn* chromosomes (Figures 2b and S2b, pink). Ingress of H3K27me3 at centromeres was consistent with previous studies showing increased H3K27me3 at chromocenters in interphase in the absence of H3K9me3^42–44^. We also detected significant increases in histone H3 serine10 phosphorylation (H3S10ph) on *Suv39h dn* metaphase chromosomes (Figure 2c, yellow) as compared to WT equivalents. Increased H3K27me3 levels and ingress of this mark at centromeric domains was also seen in mitotic chromosomes isolated from *Suv39h dn* MEFs (Figure S2c), but not on equivalent chromosomes isolated from either *Setdb1/Setdb2 double null* or *G9a/Glp double null* mutant cells (Figure S2c).

**Figure 2.**
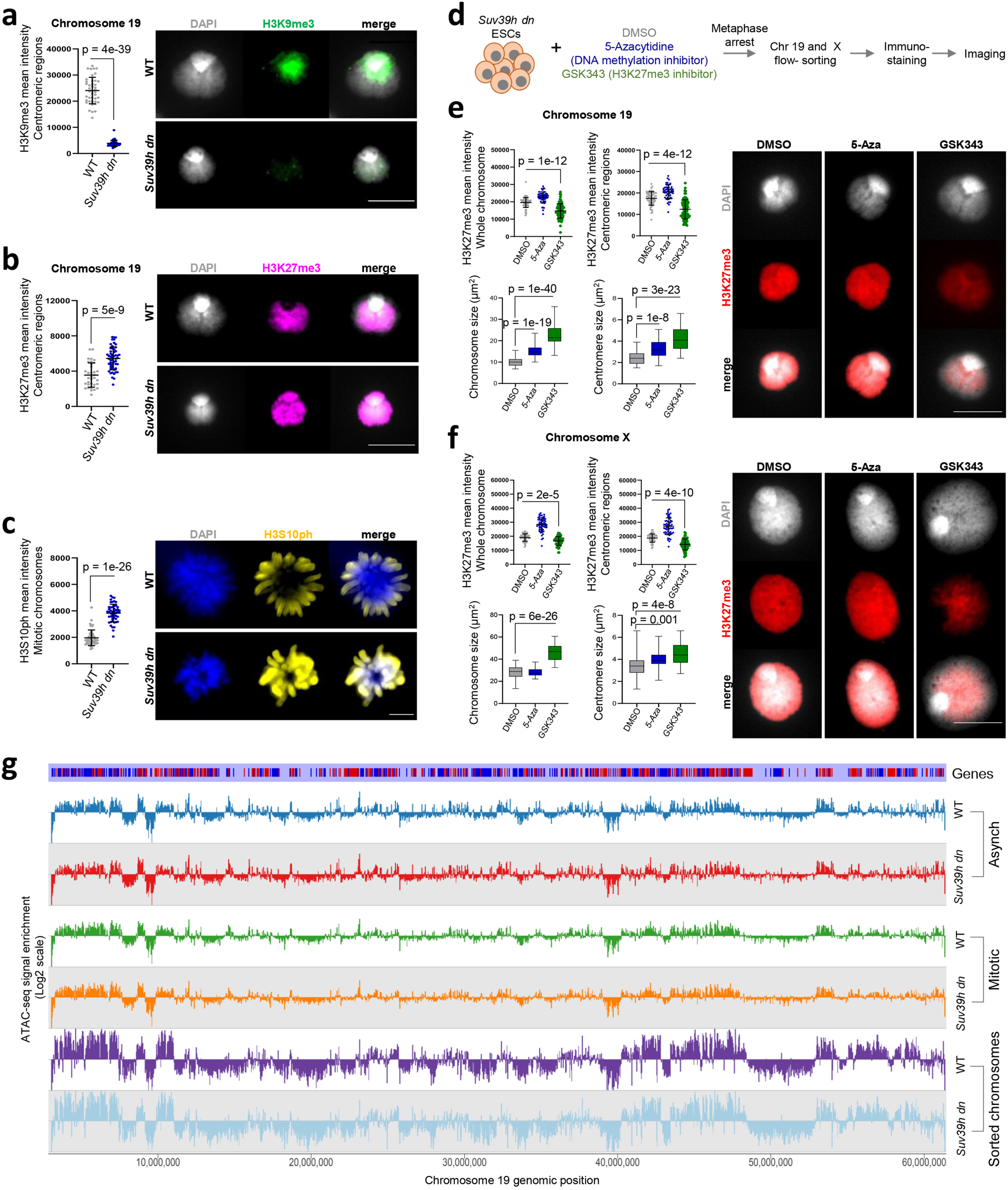
Suv39h dn mitotic chromosomes show elevated levels of H3K27me3 and H3S10ph and can be re-sized by inhibiting PRC2 activity. (a,b) Representative images (right) of immunofluorescence labelling of histone H3K9me3 (a) (green) or histone H3K27me3 (b) (pink) on chromosome 19 isolated from WT or *Suv39h dn* ESCs where DAPI counterstain is shown in light grey. Scale bars = 5 μm. Plots (left of the images) show H3K9me3 (a) or H3K27me3 (b) mean intensities, measured at centromeric regions. (c) Representative images of immunofluorescence labelling of histone H3S10ph (yellow) on WT and *Suv39h dn* metaphase-arrested ESCs, where DAPI counterstain is shown in blue. Scale bars = 4 μm. H3S10ph mean intensities were measured on mitotic chromosomes for each condition. (a-c) n = minimum 40 chromosomes analysed for each cell line over three independent experiments. P-values of statistically significant changes, measured by unpaired two tailed Student’s t-tests, are indicated. (d) Experimental strategy used to measure mitotic chromosome size of *Suv39h dn* ESCs after treatment with DNA methylation or PRC2 inhibitors (5-Aza or GSK343 respectively). (e,f) Representative images of immunofluorescence labelling of histone H3K27me3 (red) on mouse chromosome 19 (e) and chromosome X (f) isolated from *Suv39h dn* ESCs pre-treated with DMSO, 5-Aza, or GSK343. DAPI counterstain is shown in light grey. Scale bars = 5 μm. H3K27me3 mean intensities were measured at centromeric regions and whole chromosomes; mean ± SD is shown. n = minimum 50 chromosomes analysed over three independent experiments. P-values of statistically significant decreases, measured by unpaired two tailed Student’s t-tests, are indicated. Box plots show area measurements of individual chromosomes and centromeres for each condition. Minimum, lower quartile, median, upper quartile and maximum values are indicated. n = minimum 100 chromosomes analysed for each condition over three independent experiments. P-values of statistically significant increases, measured by unpaired two tailed Student’s t-tests, are indicated. (g) Chromatin accessibility profile across chromosome 19 for WT and *Suv39h dn* asynchronous and mitotic ESCs, and flow-sorted mitotic chromosomes, shown as Log2 enrichment of ATAC-seq signal.

### *Suv39h dn* mitotic chromosome size is restored by inhibiting PRC2 activity

To investigate whether enhanced compaction of *Suv39h dn* mitotic chromosomes was the result of compensatory increases in H3K27me3 (or other chromatin modifications) rather than loss of H3K9me3 *per se, Suv39h dn* ESCs were treated for 48 hours with drugs that either inhibit PRC2 activity (GSK343, an Ezh2 methyltransferase inhibitor) or that block DNA methylation (5-Azacytidine) (Figure 2d, schematic). Pre-treatment with GSK343 reduced global H3K27me3 levels on mitotic chromosomes 19 (Figure 2e, upper panel) and X (Figure 2f, upper panel), and effectively restored chromosome size and centromere compaction to that of WT equivalents (Figures 2e and 2f, lower panels). Pre-treatment of *Suv39h dn* ESCs with 5-Azacytidine (5-Aza) resulted in a smaller increase in mitotic chromosome size and centromere decompaction. In WT ESCs, treatment with either drug resulted in modest increases in mitotic chromosome size (Figure S2d). Collectively these data show that the compact structure of *Suv39h dn* mitotic chromosomes can be alleviated by inhibiting the activity of PRC2. To exclude that chromatin accessibility was grossly altered in *Suv39h dn* samples, ATAC-seq was performed on WT and *Suv39h dn* asynchronous ESCs, mitotic-arrested ESCs, as well as sorted ESC mitotic chromosomes. Global ATAC-seq profiles of *Suv39h dn* and WT chromosomes indicated that they were broadly comparable as illustrated for chromosome 19 (Figure 2g) and as shown by differential accessibility analysis (Figures S2e). This suggests that changes in chromosome-scale compaction observed in *Suv39 dn* ESCs are independent of local changes in chromatin accessibility.

### Systematic analysis of proteins bound to *Suv39h dn* mitotic chromosomes

In order to analyse the factors binding to unfixed (native) metaphase chromosomes in *Suv39h dn* and WT samples (MEFs and ESCs), we performed proteomic analyses using LC-MS/MS in which the data were analysed using the label-free quantification (LFQ) algorithm in the MaxQuant platform as detailed previously^36^. For these experiments an equivalent number of metaphase chromosomes (10^7^) in samples pre- and post-chromosome sorting were compared (schematically shown in Figure S1a) in at least three biological replicates loaded in technical duplicates on the LC-MS/MS (Figures S3a-d). This enabled discrimination of proteins that were co-enriched with mitotic chromosomes (red), were depleted (blue), or were unaltered in abundance between mitotic lysate pellets and purified chromosomes (grey) (Figures 3a and S3c). Among WT and *Suv39h dn* MEF samples, 5789 and 5784 candidate proteins were detected respectively, with a similar proportion (11-16%) showing co-enrichment with purified mitotic chromosomes (Figure 3a). Comparisons between WT and *Suv39h dn* samples revealed that many of these chromosome-bound proteins were common to both genotypes (464 of 848), although a subset of candidates (287 of 848) was not enriched in *Suv39h dn* MEF samples as compared to WT (Figure 3b). These proteins clustered in function, being associated with processes such as transcription, chromosome organisation, cell cycle, development, and nucleosome organisation (Figure 3c), in addition to H3K9 trimethylation. No obvious functional group was seen among the small group of factors (97 of 848) that were preferentially detected on *Suv39h dn* MEF mitotic chromosomes, compared to WT.

**Figure 3.**
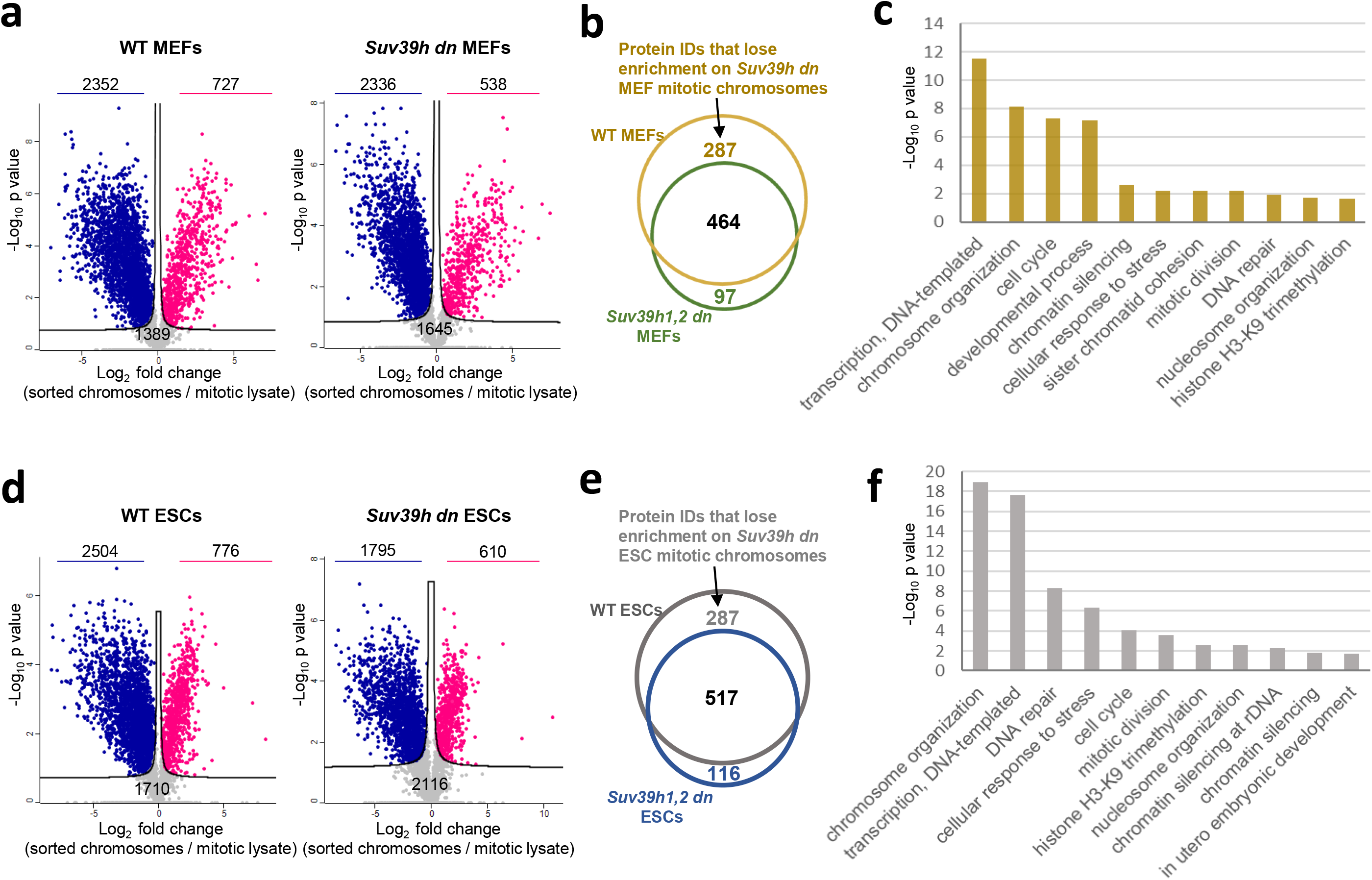
Proteomic analysis reveals a cadre of chromosome-binding factors that require H3K9me3 to remain efficiently bound during mitosis. (a) Volcano plots of proteins significantly enriched (red), depleted (blue) or not significantly changed (grey) on sorted chromosomes relative to mitotic lysate pellets for WT (left) and *Suv39h dn* (right) MEFs (unpaired two tailed Student’s t-test, permutation-based FDR < 0.05, s0 = 0.1, n = 3 independent experiments each measured in technical duplicate, see Methods for details) (Proteins were plotted as Log2 fold change LFQ intensity of sorted chromosome pellet over LFQ intensity of mitotic lysate pellet versus significance (-Log10 p) using Perseus software). The number of proteins in each category is indicated on the volcano plot. (b) Venn diagram showing the overlap of protein IDs enriched on mitotic chromosomes between WT and *Suv39h dn* MEF samples. (c) GO term analysis of proteins that lose enrichment on *Suv39h dn* mitotic MEF chromosomes compared to WT. (d-f) Proteomic comparisons as in (a-c) but for WT and *Suv39h dn* mouse ESCs.

We performed an analogous comparison in ESCs (Figures 3d-f). In WT and *Suv39h dn* ESCs, 5886 and 5624 mitotic proteins were detected respectively, with 13-15% showing a significant enrichment on mitotic chromosomes (Figure 3d). Although the majority (56%) of chromosome-bound candidates were common to both genotypes, a subset (287 of 920) was not detected in the absence of both Suv39h enzymes (Figure 3e). These candidates showed GO-term assignments that were remarkably similar to those identified previously in MEFs, encompassing chromosome organisation, transcription, DNA repair, cellular responses to stress, chromatin silencing and H3K9 trimethylation (compare Figures 3c and 3f). Taken together, these data highlight an interesting cadre of chromosome-binding proteins that apparently require H3K9me3 to remain efficiently bound in mitosis.

### Altered retention of pluripotency factors on *Suv39h dn* mitotic chromosomes

To further investigate the factors that require H3K9me3 to remain efficiently bound to mitotic chromosomes in dividing ESCs, we looked in detail at the representation of pluripotency-associated proteins in sorted mitotic chromosomes from WT and *Suv39h dn* ESC samples. Factors such as Sox2, Utf1, Zfp296, Sal4, Dppa2 and Dppa4, that we previously identified as being bound to unfixed mitotic ESC chromosomes^36^, showed an equivalent representation in control and mutant samples (Figure 4a). In the absence of Suv39h enzymes and H3K9me3 however, factors such as Esrrb (Estrogen related receptor beta, a well-characterised mitotic bookmarking factor^31, 32^), Tead4 (TEA domain transcription factor 4) and Tbx3 (T-box transcription factor) were no longer enriched on mitotic chromosomes (Figure 4a). To exclude that such deficits were simply the result of lower levels of expression by mutant cells we compared our proteomic data with previously published transcriptomic data^27^. For both genotypes there was a strong correlation between protein abundance in mitotic lysates and chromosome samples, and between protein and transcript levels (Figure S4a). However, differences in the proteome of *Suv39h dn* mitotic chromosomes, relative to WT, appeared independent of differences in the mitotic lysate or underlying transcriptome (Figure S4b). This suggests that a loss of TF retention in *Suv39h dn* mitotic chromosomes is unlikely to be explained by changes in their abundance. To explore this in more detail we examined the retention of a specific candidate, Esrrb. In WT and *Suv39h dn* ESCs, Esrrb levels were broadly comparable (Figure S4c), and proteomic data confirmed that Esrrb was equivalently represented in WT and *Suv39h dn* mitotic lysates (Figure S4d). However, sorted mitotic chromosomes from *Suv39h dn* ESCs showed an under-representation of Esrrb (Figure S4e) as compared to equivalent WT controls. To validate this result, we isolated native mitotic chromosomes 19 and X from *Suv39h dn* and WT ESCs and analysed Esrrb levels by immunofluorescence. As shown in Figure 4b, Esrrb (red) was significantly reduced in *Suv39h dn* samples as compared to WT. Interestingly, the decreased retention of Esrrb on *Suv39h dn* on mitotic chromosomes was not accompanied by loss of chromatin accessibility at specific Esrrb bookmarking sites^32^ (Figure S4f), consistent with a global similarity in ATAC-seq data derived from WT and *Suv39h dn* samples (Figures 2g and S2e).

**Figure 4.**
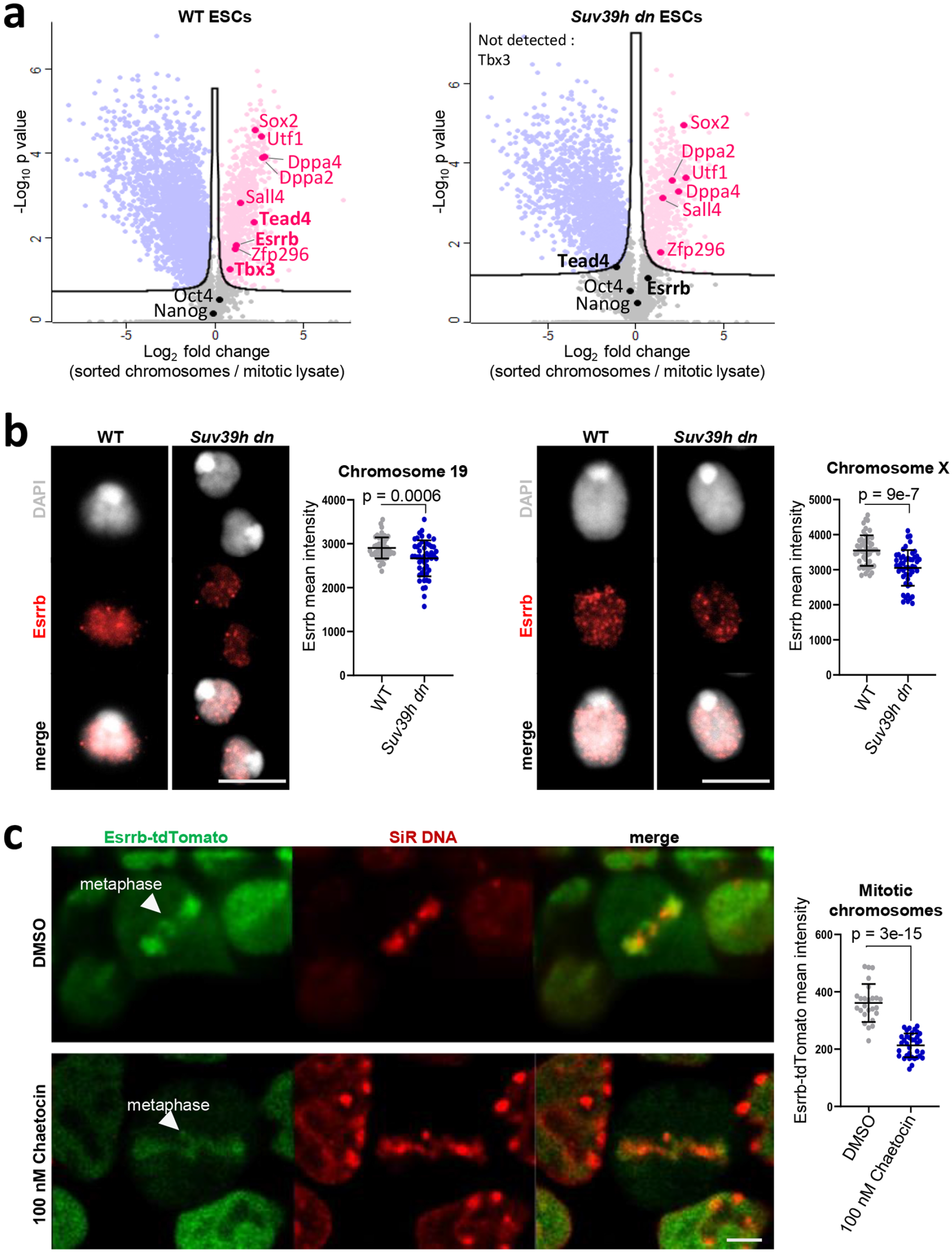
ESCs lacking Suv39h1/Suv39h2 show an altered retention of pluripotency-associated factors on mitotic chromosomes. (a) Volcano plots as in Figure 3d, highlighting pluripotency-associated factors that are enriched (red) or not significantly changed (black) on WT (left plot) or *Suv39h dn* (right plot) ESC mitotic chromosomes. (b) Esrrb immuno-labelling (red) of WT and *Suv39h dn* flow-sorted chromosomes 19 (left panel) and X (right panel), where DAPI counterstain is shown in light grey. Scale bars = 5 μm. Esrrb mean intensities were measured across individual chromosomes; mean ± SD is shown. n = minimum 50 chromosomes analysed over three independent experiments. P-values of statistically significant changes, measured by unpaired two tailed Student’s t-tests, are indicated. (c) Live-cell imaging of Esrrb-tdTomato mouse ESCs pre-treated with DMSO (upper panel) or 100 nM of Chaetocin (lower panel) cultured with SiR-DNA (red). Arrows show Esrrb localisation to mitotic chromatin, scale bars = 5 μm. Esrrb-tdTomato mean intensities on mitotic DNA (gated based on SiR-DNA signal) were quantified for each condition, mean ± SD is shown. n = 25-35 cells analysed, representative of three independent experiments. P-values of statistically significant changes, measured by unpaired two tailed Student’s t-tests, are indicated.

To test the impact of acute H3K9me3 depletion on mitotic retention of Esrrb, we used WT ESCs expressing an endogenous Esrrb-tdTomato-fusion protein^31^. Esrrb-tdTomato ESCs were examined by live cell imaging with and without addition of Chaetocin, a mycotoxin that inhibits Suv39h1 activity^45, 46^. As anticipated, Esrrb decorated metaphase chromosomes (red) in these ESCs (Figure 4c, upper panels). However following Chaetocin treatment, Esrrb binding at metaphase chromosomes was substantially reduced (Figure 4c, lower panel), although the treated cells continued to divide in culture (Figure 4c, graph right shows quantification of Esrrb intensity on mitotic DNA). Taken together these data show that both short and long-term depletion of Suv39h1/Suv39h2 activity in ESCs results in reduced retention of Esrrb on mitotic chromosomes.

### Chromosome size, centromere compaction and mitotic chromosome-bound factors are dependent on Suv39h activity

These data raise the intriguing possibility that mitotic bookmarking is sensitive to H3K9me3. As altered binding could reflect a dependency either on H3K9me3 itself, or the correct marking and function of constitutive heterochromatin domains, we asked whether other H3K9me3-associated proteins were also enriched or depleted in mitotic samples (Figure S5a). Interestingly, HP1α (or CBx5) binding was retained on *Suv39h dn* mitotic chromosomes as were the Swi/Snf chromatin re-modellers Smarcb1 and Atrx, a protein known to bind at heterochromatic repeat elements including telomeres, rDNA repeats, endogenous retroviral elements and pericentric domains^47, 48^. In contrast, components of the lysine-specific histone demethylase complex 1A (Lsd1 or Kdm1a), that are known to interact with Esrrb in trophoblast stem cells^49^, were less well retained on H3K9me3-depleted mitotic chromosomes (compare Kdm1a and Rcor1, Figure S5a). Our proteomic comparisons also revealed increases in the representation of certain factors, notably histone H1 variants, at mitotic *Suv39h dn* chromosomes, relative to WT controls (Figure S5b). This may be relevant as linker histones have been widely implicated in chromatin condensation^50–58^ and an over-representation of histone H1 could contribute to the characteristically compact state of *Suv39h dn* mitotic chromosomes.

To ask whether inefficient retention of Esrrb binding during mitosis was reversed by H3K9me3 rescue, we transfected *Suv39h dn* ESCs with Suv39h1^39^. Provision of Suv39h1 restored H3K9me3 labelling of centromeric domains, as exemplified for mitotic chromosomes 19 and X (Figure 5a), and resulted in decreased H3K27me3 across centromeric (DAPI-intense) domains (Figure 5a). Importantly, Suv39h1 transfection also rescued mitotic chromosome size and centromere compaction (Figure 5b), and restored efficient Esrrb retention by *Suv39h dn* mitotic chromosomes, assessed by proteomic profiling (Figure 5c).

**Figure 5.**
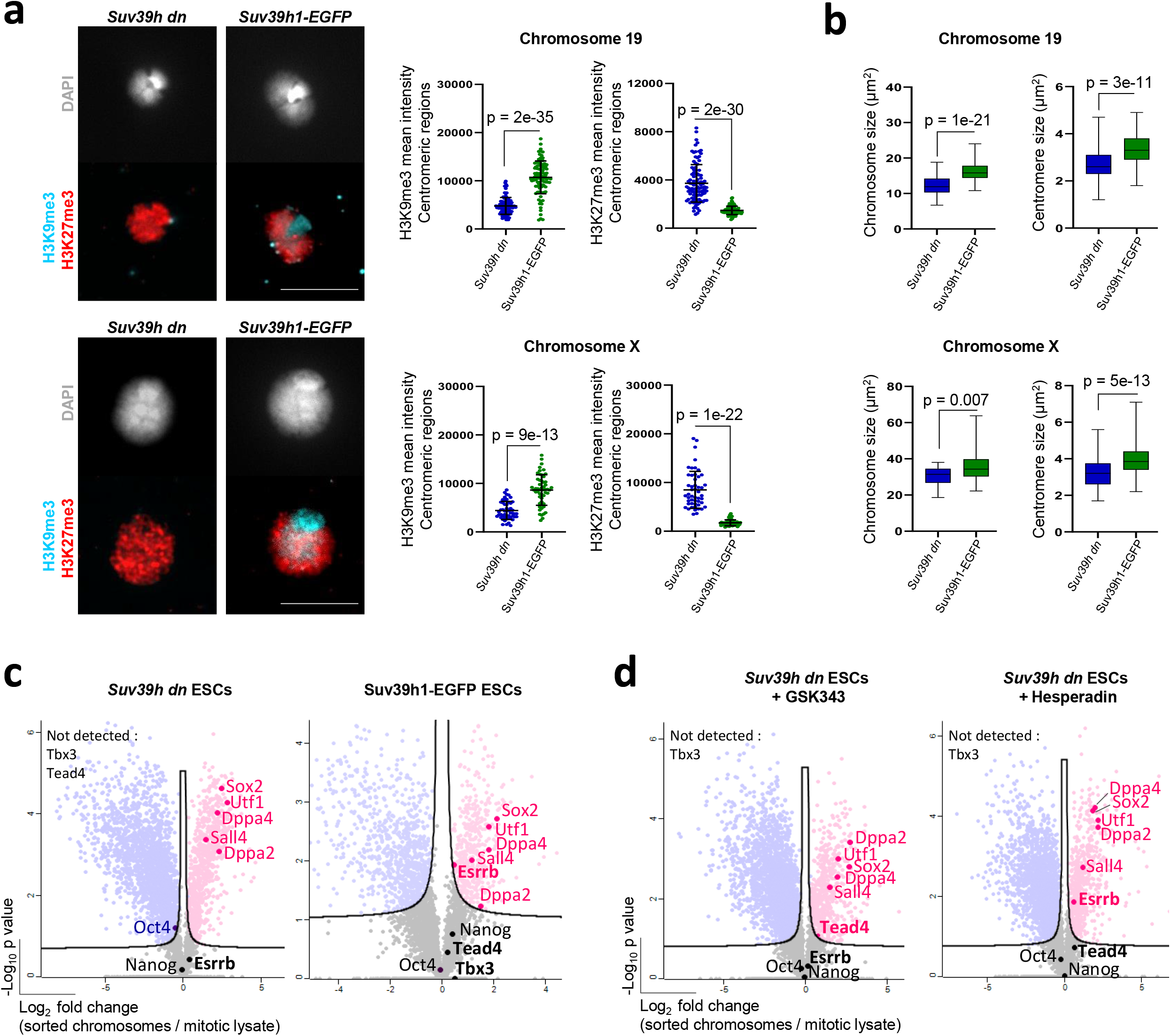
Optimal retention of mitotic bookmarking factors such as Esrrb requires H3K9me3. (a) Representative images of co-immunolabelling of histone H3K9me3 (blue) and H3K27me3 (red) on chromosomes 19 (upper panel) and X (lower panel) isolated from *Suv39h dn* ESCs or *Suv39h dn* ESCs over-expressing *Suv39h1-EGFP*. DAPI counterstain is shown in light grey. Scale bars = 5 μm. H3K9me3 and H3K27me3 mean intensities were measured at centromeric regions of chromosomes 19 and X, mean ± SD is shown. n = minimum 50 chromosomes analysed over three independent experiments. P-values of statistically significant changes, measured by unpaired two tailed Student’s t-tests, are indicated. (b) Size measurements of chromosomes 19 (upper panel) and X (lower panel) isolated from *Suv39h dn* ESCs or *Suv39h dn* ESCs over-expressing *Suv39h1-EGFP*. Box plots show area measurements of individual chromosomes and centromeres for each cell line. Minimum, lower quartile, median, upper quartile and maximum values are indicated. n = minimum 100 chromosome measurements for each cell line. P-values of statistically significant changes, measured by unpaired two tailed Student’s t-tests, are indicated. (c,d) Volcano plots highlighting pluripotency-associated factors that are enriched (red) or not significantly changed (black) on mitotic chromosomes versus mitotic lysate pellets for *Suv39h dn* ESCs or *Suv39h dn* ESCs expressing *Suv39h1-EGFP* (c), and *Suv39h dn* ESCs treated with GSK343 or *Suv39h dn* ESCs treated with Hesperadin (d). Proteins were plotted as Log2 fold change LFQ intensity of sorted chromosome pellet over LFQ intensity of mitotic lysate pellet and significance (−Log10 p) using Perseus software (unpaired two tailed Student’s t-test, permutation-based FDR < 0.05, s0 = 0.1; n = 3 independent experiments each measured in technical duplicate, see Methods for details).

### Impact of H3K9me3 loss on chromatin and chromosome-bound factors in mitosis

We have shown that H3K9me3 removal results in increased levels of H3K27me3 and H3S10ph on mitotic chromosomes. As these chromatin changes might also contribute to the altered binding of factors such as Tead4 and Esrrb in *Suv39h dn* cells, we asked whether inhibition of PRC2 activity (GSK343) or Aurora kinase b activity (Hesperadin)^59^ (to reduce H3K27me3 or H3S10ph respectively), impacts mitotic retention. Proteomic comparisons (Figure 5d) showed that Esrrb retention by *Suv39h dn* mitotic chromosomes was increased by inhibiting H3S10ph, while Tead4 retention was selectively increased upon H3K27me3 inhibition. These results reveal a clear and selective role for histone modifications in conveying mitotic bookmarking. To understand how Esrrb retention during mitosis might be impacted by the loss of H3K9me3 and altered heterochromatin structure^41^, we examined the detailed distribution of Esrrb through the cell cycle, relative to heterochromatic and euchromatic chromatin features. Using the tdTomato-Esrrb ESC line^31^, endogenous Esrrb clearly decorated chromosomes throughout all stages of mitosis (Figure S6a) and also co-localised with DAPI-stained DNA in interphase, as reported previously^31^. We show here that euchromatic and heterochromatic regions of chromosomes are labelled by Esrrb (Figures 6a and S6a) with signal also detected at DAPI-intense peri-centromeric domains of isolated mitotic chromosomes (Figure 6b). Treatment with the Su(var)3-9 inhibitor Chaetocin, substantially reduced Esrrb detection throughout mitosis (metaphase and telophase stages) and affected Esrrb distribution at euchromatin and heterochromatin domains (Figure 6c). As H3K9me3 can regulate the expression of genomic repeat elements^27^ we examined the expression of euchromatin- and heterochromatin-based repeats in *Suv39h dn* mitotic ESCs. Cot-1 RNA, that predominately contains LINE-1 and SINE repeat elements, was used to probe euchromatin repeat expression^60^ and gamma-satellite (ɣSat) RNA to probe heterochromatin-repeat expression^61^. We observed a marked increase in Cot-1 RNA in mitotic *Suv39h dn* ESCs as compared with WT controls (pink, Figure 6d), and more specifically LINE-1 expression, as confirmed by qPCR analysis (Figure S6b) and L1 ORF1 protein levels (Figure S6c). In contrast, the expression and distribution of MaSat RNA (yellow, ɣSat, Figure 6d) appeared similar in WT and *Suv39h dn* ESCs through mitosis.

**Figure 6.**
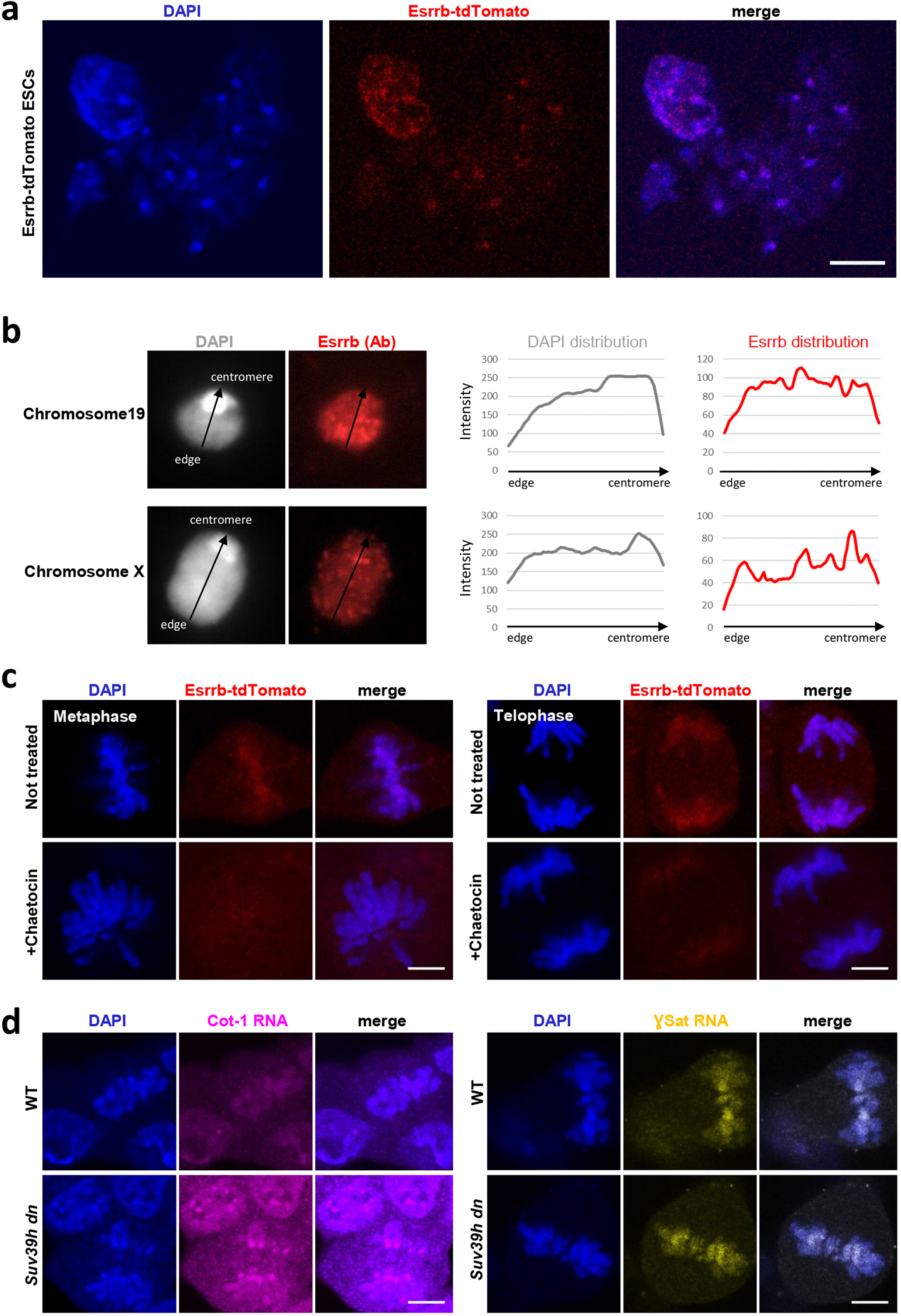
Esrrb association to euchromatin and heterochromatin regions during mitosis. (a) Representative images of Esrrb-tdTomato ESC metaphase spreads stained with DAPI (blue), scale bars = 10 μm. (b) Esrrb immuno-labelling (red) of flow-sorted chromosomes 19 (top panel) and X (lower panel), where DAPI counterstain is shown in light grey. Scale bars = 5 μm. Linescan analysis (profile plots) of Esrrb (red) and DAPI (grey) intensities across chromosome 19 and X (right panels). (c) Representative fluorescence images of Esrrb-tdTomato ESCs at metaphase (left) and telophase (right) stages with and without Chaetocin treatment. DAPI stain in blue. (d) Cot-1 (pink) and gamma-satellite repeat (yellow) RNA-FISH in WT and *Suv39h dn* ESCs, where DAPI stain is shown in blue. Scale bars = 5 μm. All the images are representative of three independent experiments.

## Discussion

This study shows that the mitotic bookmarking factor Esrrb requires H3K9me3 to be stably retained at mitotic chromosomes. Genetic ablation of Suv39h1 and Suv39h2 resulted in a reduced representation of Esrrb on native sorted mitotic chromosomes as revealed by quantitative proteomics, as well as reduced Esrrb protein detected by cellular fluorescence-based labelling. This dependency of Esrrb was supported independently by studies of acute H3K9me3 depletion targeted by pharmacological exposure to Chaetocin, and demonstrations that Esrrb representation on *Suv39h dn* mitotic chromosomes was restored by transfection of Suv39h1 and H3K9me3 rescue. Our results highlight a previously unrecognised role for H3K9me3 in retaining TFs including Esrrb, Tead4, Tbx3 and other proteins (involved in transcription, DNA repair, chromatin silencing and chromosome organisation) on condensed mitotic chromosomes. The binding of Tead4, a factor involved in Hippo signalling^62^, was diminished in *Suv39h dn* mitotic ESC chromosomes as was Tbx3, a factor expressed early in the developing ICM and implicated in regulating extraembryonic endoderm^63^ and the mitotic bookmaking factor Esrrb^31, 32^. In contrast, the pluripotency factor Sox2 that is implicated in regulating target genes in association with either Oct4 or Esrrb^64–66^, was equivalently represented on mitotic chromosomes with or without H3K9me3. Prior studies showing Esrrb and Sox2 remain bound to mitotic chromosomes^30^ in ESCs highlight an intriguing selective sensitivity of Esrrb to H3K9me3 depletion.

ATAC-seq comparison of WT and *Suv39h dn* asynchronous ESCs, mitotic cells and isolated native mitotic chromosomes revealed a surprising similarity in the profiles of mutant and control samples at a genome-wide level, as well as at Esrrb bookmarked sites (Figures 2g, S2e and S4f). This suggests that despite diminished retention of Esrrb on mitotic chromosomes lacking H3K9me3, canonical binding sites for the protein remain accessible. It is possible that alternative factors bind at these sites and that chromatin changes downstream of H3K9me3 loss negatively impact Esrrb retention at mitotic chromosomes. As an example of this we showed that increased H3S10 phosphorylation, a consequence of H3K9me3 depletion, correlates with reduced representation of Esrrb and the Esrrb-associated factors Rreb1, Tfeb^49^, and Tcf7l1. Our analyses also showed that inhibition of Aurora kinase b activity in *Suv39h dn* ESCs restored Esrrb representation on mitotic chromosomes.

To gain a broad understanding of the impact H3K9 methylation loss has during mitosis, we examined the size and epigenetic features of chromosomes isolated from ESCs and fibroblasts that lacked Suv39h1/Suv39h2, Setdb1/Setdb2 or G9a/Glp. Mitotic chromosomes from cells devoid of H3K9me3 (Suv39h1/Suv39h2 double null) were distinct in being significantly more compact than WT equivalents, with highly condensed centromeres decorated by H3K27me3 (rather than H3K9me3). These mitotic chromosomes showed a two-fold enrichment in H3S10ph and significantly increased representation of several histone H1 variants as determined by proteomic analysis (histones H1.1, H1.2, H1.3, H1.4 and H1.5, Figure S5b).

Increased levels of histone H1 variants, H3S10ph and H3K27me3 have independently been shown to result in enhanced chromosome compaction in other settings^36, 50–52, 67–69^ and we show here that inhibition of H3K27me3 activity can reverse the compaction of *Suv39h dn* mitotic chromosomes. Taken together these results contribute to an extensive catalogue of structural and organisational defects that have previously been described in interphase mammalian cells upon H3K9me3 withdrawal^70, 71^. This includes altered heterochromatin organisation, extended telomere length, DNA repair pathway activation, repeat element re-expression and genomic instability that manifests as H3K9me3-deficient cells transit mitotic or meiotic division^13, 41^. Our data, that are focused on mitotic events, reveal that H3K9me3 is also required to sustain binding of a rich cadre of proteins to mitotic chromosomes. These proteins collectively influence DNA-templated transcription, chromatin silencing, DNA repair, cellular stress and chromosome organisation, processes that are likely to be particularly important during and immediately after cell division. In this regard it is worth noting that Suv39h1/Suv39h2 deficient mice show severely impaired viability, aneuploidy with increased tumour incidence, and also fail to generate mature functional sperm^13^.

We have shown that cells lacking H3K9me3 methyltransferase enzymes Suv39h1 and Suv39h2 display consistent changes in the repertoire of factors that stably bind to mitotic chromosomes, that cannot be solely attributed to underlying changes in gene/protein expression. This observation provides important primary evidence that mitotic bookmarking is sensitive to chromatin context. Our analysis of Suv39h1/Suv39h2 mutant fibroblasts and ESCs reveals reduced representation of several TFs on H3K9me3-depleted mitotic chromosomes, in addition to proteins that are involved in DNA repair, nucleosome organisation, chromatin silencing and chromosome organisation. It is interesting to speculate that these groups of proteins might be particularly relevant in protecting the genetic and epigenetic state of daughter cells as these inherit a cargo of chromosome-bound proteins following cytokinesis, and before de-novo gene expression begins early in G1. In this setting both WT and *Suv39h dn* mitotic chromosomes showed an enrichment in core PRC1, PRC2 components and DNA methylation machinery, relative to mitotic lysates. In addition, although retention of certain TFs was impaired following H3K9me3 loss, many pluripotency-associated factors such as Dppa2, Dppa4, Mpp8 and Sox2 remained chromosome-associated in Suv39h1/Suv39h2 mutants through mitosis. Dppa2, Dppa4 are small heterodimerising nuclear proteins that are known to regulate zygotic genome activation. In ESCs these proteins have also been shown to be critical for maintaining the functionally primed state of a subset of bivalent genes in ESCs, protecting them from *de novo* DNA methylation and irreversible silencing^72, 73^. M-Phase Phosphoprotein 8 (Mpp8 or Mphosph8) is an essential player in safeguarding ground-state pluripotency and stem cell renewal, through interactions with a plethora of epigenetic silencing proteins including Dnmt3a, Setdb1 and Sirt1. Interestingly Mpp8 has also been implicated in repressing LINE1 and L1ORF2 expression, in conjunction with the human silencing hub (HUSH) complex^74^. While we do not yet know the basis of mitotic factor sensitivity or resistance to H3K9me3 loss, it is possible that chromosome-bound Dppa2, Dppa4 and Mpp8 could act as safeguards, protecting the epigenetic fidelity and pluripotent state of newly established daughter cells, particularly in the context of *de novo* DNA methylation conveyed by chromosome-bound Dnmt3a. Our analyses also showed that ESCs that lack Suv39h1/Suv39h2 over-express LINE1 and L1ORF2, consistent with prior studies^27^, and we detected an increased representation of L1OF2 protein on H3K9me3-depleted mitotic chromosomes (Figures S6B, S6C). As H3K9me3 is not strictly required for Mpp8-mediated repression of LINE1 elements^74^ it is likely that the overexpression of LINE1 elements in mutant Suv39h1h2-null cells may stem from failures to recruit and maintain Setdb1 and the HUSH complex at appropriate targets.

This study asks whether repressive chromatin influences genomic bookmarking in mitosis. This important question was addressed using an approach that enables unfixed ‘native’ mitotic chromosomes to be directly isolated from dividing cells^36^. Examining purified mitotic chromosomes from cells either at different stages of development, or from cells that lack specific chromatin features, it was possible to compile a comprehensive and quantitative catalogue of proteins that remained bound to mitotic chromosomes in the absence of Suv39h1/Suv39h2-mediated H3K9me3. These data revealed an unexpected role for repressive chromatin in maintaining the efficient binding of selected proteins to mitotic chromosomes. We therefore show that in addition to the important role of H3K9me3 in regulating gene and chromosome function during interphase, H3K9me is important for mitotic chromosome structure and moreover, for the correct genomic bookmarking of mitotic chromosomes.

## Materials and Methods

### Cells

Mouse ESCs used in this study were WT ESC (WT26), *Suv39h dn* (DN57)^26^, *Suv39h1-EGFP* (*Suv39h dn* ESCs overexpressing full length Suv39h1)^39^, Esrrb-tdTomato^31^ (gift from Nicola Festuccia). All the ESC lines were grown on 0.1% gelatin (G1993, Sigma-Aldrich) coated dishes (feeder-free) in KO-DMEM medium (Gibco, 10829018) supplemented with 15% FBS (Gibco, 10270106), 2 mM L-glutamine (25030081, Gibco), 1x MEM non-essential amino acids (Gibco, 11140035), 100 μM 2-mercaptoethanol (31350010, Gibco), antibiotics (10 μg/ml Penicillin and Streptomycin, 15140122, Gibco) and 1000 U/ml of leukaemia inhibitory factor (LIF). Mouse Embryonic Fibroblasts (MEFs) used in this study were WT (W8), *Suv39h dn* (D5)^13^, WT (clone Eset25 control), *Suv39h1/2 −/−* (CRISPR clone B1), *Setdb1/2 −/−* (CRISPR clone A4+OHT) and *G9a/Glp −/−* (CRISPR clone H7)^41^. MEFs were maintained in high-glucose DMEM supplemented with 10% FBS, 2 mM L-glutamine, antibiotics (10 μg/ml Penicillin and Streptomycin), 50 μM 2-mercaptoethanol, 1x non-essential amino acids and sodium pyruvate. For A4 cells, Setdb1 deletion was induced by plating cells in 2 μM 4-OHT for 2 days, followed by 2 days recovery in complete MEF medium without tamoxifen.

### Drug treatments

Inhibitors used in this study were Ezh2 inhibitor GSK343 (SML0766, Sigma-Aldrich), DNA methylation inhibitor 5-Azacytidine (A2385, Sigma-Aldrich), Suv39h1 inhibitor Chaetocin (GR349-0200, Enzo-LifeScience) and Aurkb inhibitor Hesperadin (375680, Sigma-Aldrich). Stock solutions were prepared to 10 mM GSK343, 100 mM 5-Aza, 10 mM Chaetocin, and 10 mM Hesperadin in DMSO. 24 h after passaging the ESC lines, cells were treated in 10 cm dishes with 1 μM GSK343, or 100 nM 5-Aza, or 100 nM Chaetocin, or 100 nM of Hesperadin) for 24 h at 37°C.

### Mitotic chromosome preparation and flow-sorting

Mitotic chromosomes were prepared using a polyamine-based method and flow-sorted as described previously^36, 75^. ESCs and MEFs were arrested in metaphase using 0.1 μg/ml demecolcine (D1925, Sigma-Aldrich) for 6 h at 37°C. Metaphase-arrested cells were collected by mitotic shake off and 1x PBS washes. Approximately 10 million mitotic cells were centrifuged at 289 g for 5 min at room temperature (RT). Mitotic cell pellets were gently resuspended in 10 ml of hypotonic solution (75 mM KCl, 10 mM MgSO_4_, 0.5 mM Spermidine trihydrochloride (S2501, Sigma-Aldrich) 0.2 mM Spermine tetrahydrochloride (S2876, Sigma-Aldrich); adjusted to pH 8 with 0.25 M NaOH) for 20 min at RT. Swollen cells were centrifuged at 300 g for 5 min at RT, resuspended in 3 ml of freshly prepared ice-cold polyamine isolation buffer (15 mM Tris-HCl, 2 mM EDTA, 0.5 mM EGTA, 80 mM KCl, 3 mM DTT, 0.25% Triton X-100, 0.2 mM Spermine tetrahydrochloride and 0.5 mM Spermidine trihydrochloride; pH 7.5), and incubated in the polyamine buffer for 15 min on ice. Mitotic chromosomes were released by vortexing (maximum speed) for 30 sec and syringing through a 23-gauge needle using a 1 ml syringe. Chromosome suspensions were then centrifuged for 2 min at 200 g at RT. The supernatant containing mitotic chromosomes was filtered using a 20 μm mesh CellTrics filter (04-0042-2315, Sysmex) into a 15 ml Falcon tube. Mitotic chromosome preparations were stained with 5 μg/ml Hoechst 33258 (94403, Sigma-Aldrich), 50 μg/ml Chromomycin A3 (C2659, Sigma-Aldrich) and 10 mM MgSO_4_, overnight at 4°C. 1 h prior to FACS sorting, 10 mM sodium citrate and 25 mM sodium sulfite were added to chromosome suspensions. Chromosomes were analysed and sorted by flow cytometry using a Becton Dickinson Influx equipped with spatially separated lasers. Hoechst 33258 was excited using a (Spectra Physics Vanguard, air cooled) 355 nm laser with a power output of 350 mW. Hoechst 33258 fluorescence was collected using a 400 nm long pass filter in combination with a 500 nm short pass filter. Chromomycin A3 was excited using a (Coherent Genesis, water cooled) 460 nm laser with a power output of 500 mW. Chromomycin A3 fluorescence was collected using a 500 nm long pass filter in combination with a 600 nm short pass filter. Forward scatter was measured using a (Coherent Sapphire) 488 nm laser with a power output of 200 mW and this was used as the trigger signal for data collection. Chromosomes were sorted at an event rate of 20000 per second. A 70-micron nozzle tip was used along with a drop drive frequency set to ~96 KHz and the sheath pressure was set to 65 PSI.

### Immunofluorescence on flow-sorted chromosomes

Flow-sorted chromosomes (100,000 chromosomes 19 or X) were cytospun onto Poly-L-lysine coated slides by cytocentrifugation (Cytospin3, Shandon) at 1,300 rpm for 10 min at RT. Chromosome samples (unfixed) were incubated with blocking buffer (5% normal goat serum in a buffer containing 10 mM HEPES, 2 mM MgCl_2_, 100 mM KCl and 5 mM EGTA) for 30 min at RT, and incubated overnight at 4°C in a humid chamber with primary antibodies to Cenpa (2040S, Cell Signalling), H3K9me3 (07-523 or 07-442, Millipore), H3K27me3 (ab6002, Abcam or 07-449, Millipore), Esrrb (PP-H6705-00, Perseus Proteomics). All antibodies were diluted 1/200 in the blocking buffer. Chromosomes were then incubated with appropriate secondary antibodies (anti-mouse-Alexa488 (A11001, Invitrogen), anti-rabbit-Alexa488 (A11008, Invitrogen), or anti-mouse-A566 (A11031, Invitrogen)) for 1 h at RT. All secondary antibodies were diluted 1/400 in the blocking buffer. Immuno-stained chromosomes were mounted in Vectashield mounting medium containing DAPI (H-1200, Vector Laboratories). Wide-field epi-fluorescence microscopy was performed on an Olympus IX70 inverted microscope using a UPlanApo 100x/1.35 Oil Objective lens.

### Immunofluorescence on cells

ESCs were cultured on gelatin-coated glass coverslips, cells were fixed with 1% paraformaldehyde for 10 min at RT, crosslinked with 2 mM disuccinimidyl glutarate (DSG) for 50 min at RT as previously described^32^. Cells were then permeabilized with 0.1% Triton X-100 for 15 min at RT and blocked with 1% bovine serum albumin and 5% goat serum (Sigma). cells were incubated with primary antibodies Esrrb (PP-H6705-00, Perseus Proteomics) or H3S10ph (ab5176, Abcam) at 4 °C overnight. After 3 washes with 1x PBS 0.05% Triton, cells were labelled with Alexa Fluor-conjugated secondary antibodies and mounted in Vectashield mounting medium containing DAPI.

### Chromosome size measurement and histone modification quantification

After flow-sorting and cytospinning, chromosome 19 and chromosome X were stained with anti-Cenpa or co-stained with anti-H3K9me3 and anti-H3K27me3 as described above. Immuno-stained chromosomes were 3D-imaged using wide-field epi-fluorescence microscopy performed on an Olympus IX70 inverted microscope using a UPlanApo 100x/1.35 Oil Objective lens. Acquired images were opened and analysed on Fiji^76^. Chromosome and centromere size measurements were assessed using a custom script in Fiji to estimate whole chromosome (total DAPI) and centromere (DAPI high) areas. After selection of centromere and whole chromosome areas, H3K27me3 and H3K9me3 maximum intensity projections of z-planes were quantified for both centromere and whole chromosome areas of each individual chromosome.

### Metaphase spreads

WT and *Suv39h dn* ESCs (80% confluency) were arrested in metaphase using 0.1 μg/ml demecolcine solution. Cells were then trypsinized, centrifuged, and re-suspended in hypotonic solution (40 mM KCl, 0.5 mM EDTA, 20 mM HEPES, pH to 7.4 using NaOH, pre-warmed to 37°C) for 25 min at 37°C. Nuclei were pelleted (8 min at 500 g) and supernatant removed (apart from a small drop to re-suspend the pellet) prior to addition of 3:1 MeOH:glacial acetic acid (both Fisher Chemical) fixative (made fresh and pre-cooled to −20°C) to the top of the tube. The tubes were stored at −20°C overnight. The next day nuclei were pelleted (8 min at 500 g) and washed in fresh 3:1 MeOH:glacial acetic acid three times before preparation of metaphase spreads. To prepare chromosome spreads nuclei were pelleted and re-suspended in a small volume of fixative (to a pale grey solution). A 20 μl drop of 45% Acetic Acid in water was pipetted onto a glass Twinfrost microscope slide and 23 μl of spread mixture dropped onto it, tilting the slide to spread the nuclei. The slides were air-dried and stored dry at room temperature. XCyting Mouse Chromosome Painting Probes to chromosomes X and 19 (Metasystems Probes) were used alone or together with mouse ɣ-satellite probes (DNA a gift from Niall Dillon) directly labelled with FluoroRed (Amersham Life Science RPN2122) by nick translation, to detect chromosomes X or 19 with pericentromeric DNA. Metaphase chromosome painting was performed according to the protocol supplied by Metasystems Probes and mounted in Vectashield containing DAPI. Leica SPII confocal microscope was used for imaging.

### ATAC-seq

ATAC-seq^77^ was performed in duplicate on asynchronous and mitotic WT and *Suv39h dn* ESCs and on purified mitotic chromosomes, as previously described^36^. Briefly, nuclei were obtained from asynchronous ESCs (5 x 10^4^) according to the Omni-ATAC-seq protocol^78^ by resuspending in 50 μl of ATAC-resuspension buffer (10 mM Tris–HCl, pH 7.4; 10 mM NaCl; 3 mM MgCl_2_; 0.1% Igepal CA-630; 0.1% Tween-20; 0.01% Digitonin) and incubating on ice for 3 min. After adding 1 ml of ATAC-resuspension buffer without Igepal and Digitonin, nuclei were pelleted at 500 *g* (10 min at 4°C). To perform ATAC-seq, nuclei, mitotic cells (5 x 10^4^) or purified chromosomes (2 x 10^6^ pelleted at 10,000g for 10min at 4°C) were resuspended in 50 μl transposase mixture (25 μl Illumina TD buffer, 22.5 μl H_2_O, and 2.5 μl Illumina TDE1 transposase) and incubated at 37°C for 30 min with shaking at 1,000 rpm. DNA was purified (Qiagen MinElute kit) and amplified with seven cycles of PCR (NEBNext High Fidelity master mix, primers shown in Supplemental Table 1)^77^. Libraries were purified twice with Ampure XP beads (Beckman Coulter), including removal of large fragments with 0.5x beads. Libraries were assessed by Qubit, Bioanalyzer and KAPA Library Quantification (Roche) before Illumina NextSeq500 sequencing (75 bp, paired end). Data were processed using RTA (v2.11.3, default settings) and reads were demultiplexed with bcl2fastq2 (v2.20.0; allowing 0 mismatches). Reads were trimmed with Trim Galore! (trim_galore_v0.4.4; --trim-n --paired; www.bioinformatics.babraham.ac.uk/projects/trim_galore) and aligned to UCSC mm10 using bowtie2 (bowtie2/2.2.9; -p8 -t -very-sensitive -X 2000)^79^. Aligned bam files were sorted with Samtools (v1.2)^80^, and bam files from technical replicate sequencing runs were merged. Duplicate reads were marked with Picardtools MarkDuplicates (v1.90; http://broadinstitute.github.io/picard/). The bamQC function from R/Bioconductor package ATACseqQC^81^ was used to keep properly mapped paired-end reads, and remove duplicates and mitochondrial alignments. Chromosome-wide accessibility profiles were generated from the resulting bam files in Seqmonk (v1.48.0; www.bioinformatics.babraham.ac.uk/projects/seqmonk) by expressing the number of transposase insertions (5’ read ends, offset by +4 bp/−5 bp) in each 25 kb window, relative to the genome-wide average. Bam files were further processed with deeptools^82^ to shift and extract nucleosome-free fragments (<100 bp). Peaks were called from nucleosome-free fragments with MACS2 (-f BAMPE)^83^. Peaks overlapping regions in the ENCODE blacklist v2^84^ were removed. For downstream analysis, we separately defined consensus peaks for cells (asynchronous and mitotic) and purified chromosomes. To generate each consensus list, we defined non-redundant peaks using the reduce function from R/Bioconductor package GenomicRanges^85^ and only kept peaks appearing in at least two samples. Peak-based read counts were obtained with the featureCounts function from R/Bioconductor package Rsubread^86, 87^ and normalised using the calcNormFactors function (method = “TMM”) in R/Bioconductor package edgeR (v3.28.1)^88, 89^. Differential accessibility analysis was performed with R/Bioconductor package limma (v3.42.2)^90^, after applying the voom function^91^ to estimate the mean-variance relationship within the count-based datasets. For accessibility trend plots, insertion sites were extended ±25 bp to smooth signal and plotted as the average relative distribution across 2 kb windows centred on Esrrb peak summits. Esrrb peak locations and bookmarking status were taken from^32^, and coordinates were converted to mm10 using the UCSC (genome.ucsc.edu) LiftOver tool.

### Proteomics

Proteomics was performed on mitotic lysate pellets and sorted chromosomes (equivalent number, 10 million) isolated from WT ESCs, *Suv39h dn* ESCs, *Suv39h dn* ESCs treated with GSK343 or Hesperadin, *Suv39h1-EGFP* ESCs, WT MEFs, *Suv39h dn* MEFs using a method previously described^36^. Both flow-sorted mitotic chromosomes and mitotic lysates were pelleted by centrifugation (14,000 g, 15 min, 4°C). Supernatants were removed and the pellets were digested with trypsin by an in-Stage Tip digestion protocol^92^ using commercially available iST Kits (Preomics, Martinsried, Germany) according to the manufacturer’s recommendations. Protein digests were analysed by liquid chromatography-tandem mass spectrometry (LC-MS/MS) using a data-dependent acquisition method with a 50 cm EasySpray column at a flow rate of 250 nl/min coupled to a Q-Exactive HF-X mass spectrometer, as described previously^36^. Experiments were carried out in biological triplicate and digests were analysed by LC-MS/MS as technical duplicates. Raw data obtained were analysed using MaxQuant^93, 94^ with the in-built Label-Free Quantification algorithm^95^. Further statistical analysis, as well as data visualisation were performed using the Perseus software platform^96^.

### Comparison of transcript abundance with proteomics

Raw poly-A RNA-sequencing data from WT and *Suv39h dn* ESCs were obtained from the NCBI Sequence Read Archive (SRA, GSE57092)^27^. Reads were aligned to the UCSC mm10 genome using the STAR aligner (v2.7.7a)^97^ based on the Ensembl genome annotation (v2.7.7)^98^. Gene-based read counts were obtained with STAR and normalised by calculating Transcripts Per Million (TPM). Differential expression analysis was performed using DESeq2 (v1.30.1)^99^ to generate log2 fold changes. Gene symbols were obtained for transcripts using Ensembl BioMart (https://www.ensembl.org/biomart/martview) and used to match transcriptomics and proteomics data. Ambiguous gene symbols were removed from the proteomics data. In total, 5664 genes were mapped between proteomics and transcriptomics datasets for correlation analyses.

### Live cell imaging

48 h prior to imaging, Esrrb-tdTomato ESCs were grown in ESC medium in Ibidi μ-Sildes 8 Well (Ibidi, 80826), pre-coated with 0.1% gelatin. Next day, cells were treated with 100 nM of Chaetocin or with DMSO in fresh ESC medium for 24 h. Approximately 1 h prior to live cell imaging, cells were switched to phenol red-free medium (31053-028, Gibco) containing 15% FBS, 1x non-essential amino acids and sodium pyruvate, 2 mM L-glutamine, 100 μM 2-mercaptoethanol, antibiotics, and LIF, buffered with 20 mM HEPES. Cells were incubated with 1 μM SiR-DNA (SC007, SpiroChrome) 30 min before imaging. Time-lapse images were acquired on an Olympus IX70 inverted widefield microscope using a UPlanApo 100x/1.35 Oil Objective lens and an environmental chamber kept at 37°C with a 5% CO2 supply. Z-stacks were collected every 120 sec with a step-size of 2 μm. Images were deconvolved with Huygens Professional version 19.10 (Scientific Volume Imaging, The Netherlands, http://svi.nl), using the CMLE algorithm. Esrrb-tdTomato fluorescence signal was measured on maximum intensity projections of z-planes for both interphase nuclei and metaphase chromosomes (based on SiR-DNA signal).

### Probe labelling

1 μg of mouse Cot-1 (18440016, Invitrogen) or gamma satellite DNA (a kind gift from Niall Dillon) was labelled with either Cy3-dUTP or Cy5-dUTP (PA53022 or PA55022 respectively, Cytiva or Sigma-Aldrich) using a Prime-It Flour Fluorescence labelling kit (300380 version B, Agilent Technologies) according to manufacturer’s instructions. Briefly, the DNA template was annealed with random 9-mer primers at 95°C for 5 min, then the RNA extended using exonuclease-free Klenow polymerase for 30 min at 37°C. The reaction was stopped and the probe stored at 4°C.

### RNA-FISH

Cells were grown on coverslips. Cells were washed with PBS, fixed using 2.6% formaldehyde in PBS for 10 min then permeabilized using 0.4% Triton X-100 in PBS for 5 min on ice. Cells were washed with PBS and dehydrated using an ethanol series (3 min in 70%, 80%, 95% and 100% EtOH). 2.5 μl of RNA probe in 12 μl of hybridization buffer (1 ml 50% Denhardt’s solution, 200 μl SSC containing 20% dextran sulphate, 800 μl formamide) was used per 22 x 22 mm coverslip. Probe was denatured at 75°C for 8 min, then placed on ice for 1 - 2 min.

Cells were inverted onto the probe on a slide, sealed with rubber cement and placed in a humid chamber at 37°C overnight (at least 16 h). Following hybridization, the rubber cement was removed and cells washed 3 x 3 min with 2x SSC containing 50% formamide at 42°C followed by 3 x 3 min washes with 2x SSC at 42°C, and briefly rinsed in water before mounting in Vector Shield containing DAPI.

### Quantitative RT-PCR

Total RNA was extracted from asynchronous or metaphase-arrested WT and *Suv39h dn* ESCs (500.000 cells) using RNeasy mini Kit (74106, Qiagen). Extracted RNA was treated twice with Turbo DNA free (AM1907, Invitrogen) according to the manufacturer’s instructions and reverse transcribed using SuperScript III (18080085, Invitrogen) and random primers (Invitrogen). Real time PCR was performed in technical triplicate for each sample, using SYBR Green PCR Master Mix (204145, Qiagen) on a real time PCR machine (Bio-Rad CFX system). Values were normalized to β-actin and Gapdh. Primer sequences used in this study are listed in supplemental table 2.

### Western blot

WT and *Suv39h dn* asynchronous ESCs (1 million) were pelleted and resuspended in 100 μl of cold RIPA buffer (50 mM Tris-HCl pH 8.8, 150 mM NaCl, 1% Triton X-100, 0.5% sodium deoxycholate, 0.1% SDS, 1 mM EDTA, 3 mM MgCl2), supplemented with 1× protease inhibitor cocktail (11873580001, Roche) and 1.25 U/μl Benzonase (E1014, Sigma). Samples were incubated for 20 min at RT, mixed with 100 μl of 2× Laemmli sample buffer (65.8 mM Tris-HCl pH 6.8, 2.2% SDS, 22.2% glycerol, 0.01% bromophenol blue and 710 mM 2-mercaptoethanol), and denatured at 95 °C for 10 min. Western blots were performed according to standard procedures using Immobilon Block-FL (WBAVDFL01, Millipore) as fluorescent blocker, and near infra-red detection was carried out using the LI-COR detection system. The following antibodies and dilutions were used: anti-Esrrb (PP-H6705-00, Perseus Proteomics, 1:1000) and anti-Histone H3 (61476, Active Motif, 1:5000).

## Acknowledgments

We thank the LMS/NIHR Imperial Biomedical Research Centre Flow Cytometry Facility, as well as the LMS microscopy facility and the LMS Genomics and LMS Bioinformatics facilities for support. We are grateful to N. Festuccia and P. Navarro for providing Esrrb-tdTomato ESC line. We thank N. Dillon for the mouse gamma satellite DNA plasmids. Research in the laboratory of T. Jenuwein is supported by the Max Planck Society and by additional funds from the German Research Foundation (DFG) within the CRC992 consortium ‘MEDEP’.

This work was funded by core support from the Medical Research Council UK to the London Institute of Medical Sciences.

