## Supplemental Information for "Loss of H3K9 tri-methylation alters chromosome compaction and transcription factor retention during mitosis"

Djeghloul, D., et al.

Supplemental Figure S1
Supplemental Figure S2
Supplemental Figure S3
Supplemental Figure S4
Supplemental Figure S5
Supplemental Figure S6
Supplemental table 1
Supplemental table 2
Supplemental references

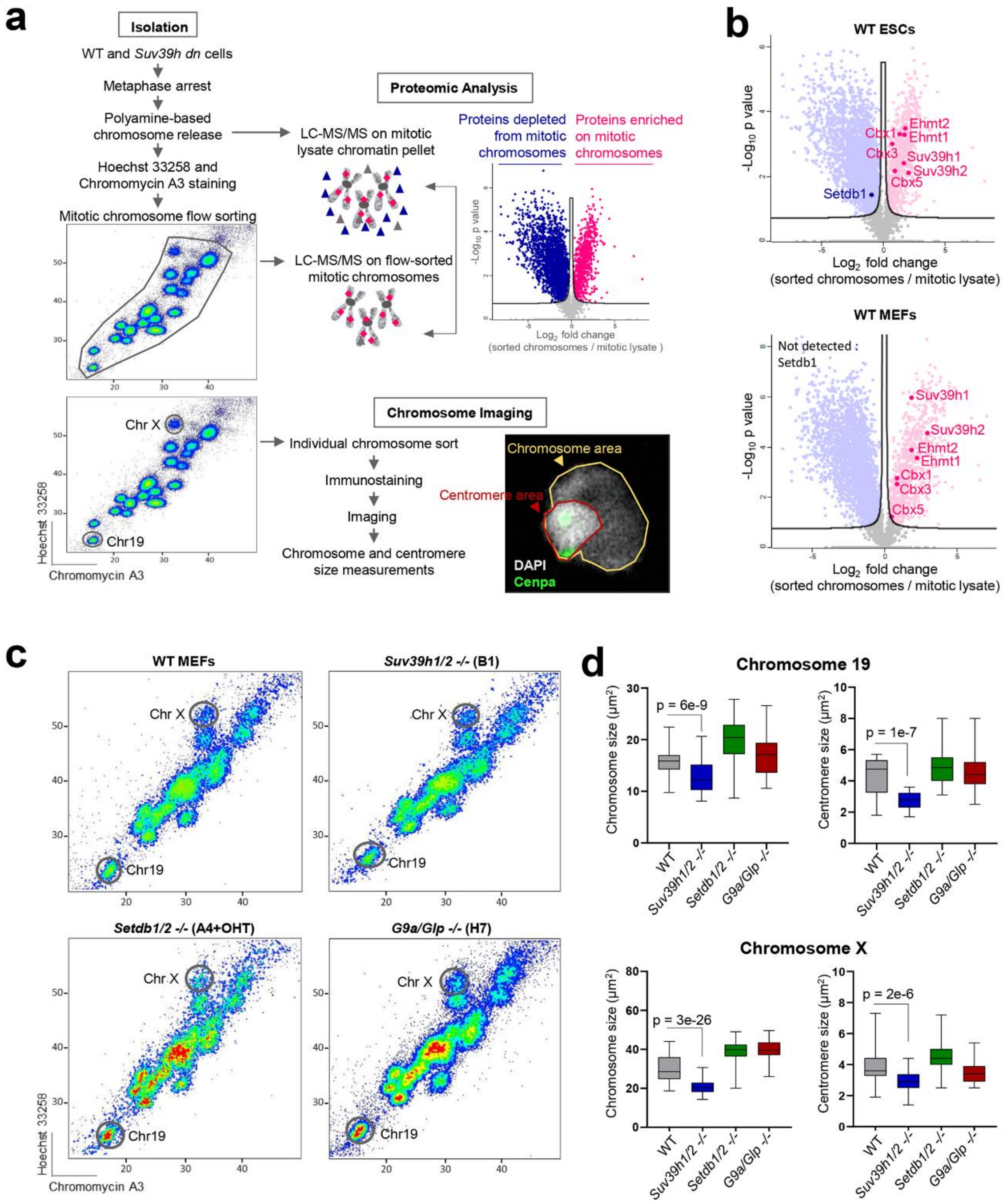


**Figure S1:** (a) Scheme of experimental strategy used to isolate native metaphase chromosomes from WT and *Suv39h dn* cells, and identify proteins bound to mitotic chromatin in each cell line. Hoechst 33258 and Chromomycin A3 bivariate karyotype was assessed by flow cytometry and the gates used to sort all chromosomes, chromosome 19, or the X chromosome are indicated. Proteomic analysis was performed using LC-MS/MS on total mitotic cell lysate pellet, or on flow-purified chromosomes, to identify proteins enriched on metaphase chromosomes. Chromosome 19 and X size measurements were performed using Fiji/imageJ software to estimate chromosome (total DAPI) and centromere (DAPI high) areas, as indicated. (b) Volcano plots of proteins detected as being significantly enriched (red), depleted (blue) or not significantly enriched (grey) on sorted chromosomes relative to mitotic lysate pellet for WT ESCs (upper plot) or WT MEFs (lower plot). H3K9 KMTs and HP1 proteins are highlighted on the volcano plots. Statistical analysis was performed using unpaired two tailed Student's t-test, permutation-based FDR < 0.05, n = 3 independent experiments each measured in duplicate, see Methods for details). Proteins were plotted as Log2 fold change (LFQ intensity of sorted chromosome pellet /LFQ intensity of mitotic lysate pellet) and significance (-Log10 p) using Perseus software. (c) Flow karyotypes of mitotic chromosomes isolated from WT, *Suv39h1/2 -/-* (B1), *Setdb1/2 -/-* (A4+OHT), or *G9a/Glp -/-* (H7) MEFs. Gates used to isolate chromosomes 19 and X from each cell line are indicated. (d) Chromosome 19 (upper panel) and X (lower panel) size measurements from WT, *Suv39h1/2 -/-* (B1), *Setdb1/2 -/-* (A4+OHT), or *G9a/Glp -/-* (H7) MEFs. Box plots show area measurements of individual chromosomes and centromeres for each MEF line. Minimum, lower quartile, median, upper quartile and maximum values are indicated. n = minimum 100 chromosomes analysed for each condition over three independent experiments. P-values of statistically significant decreases, measured by unpaired two tailed Student's t-tests, are indicated.

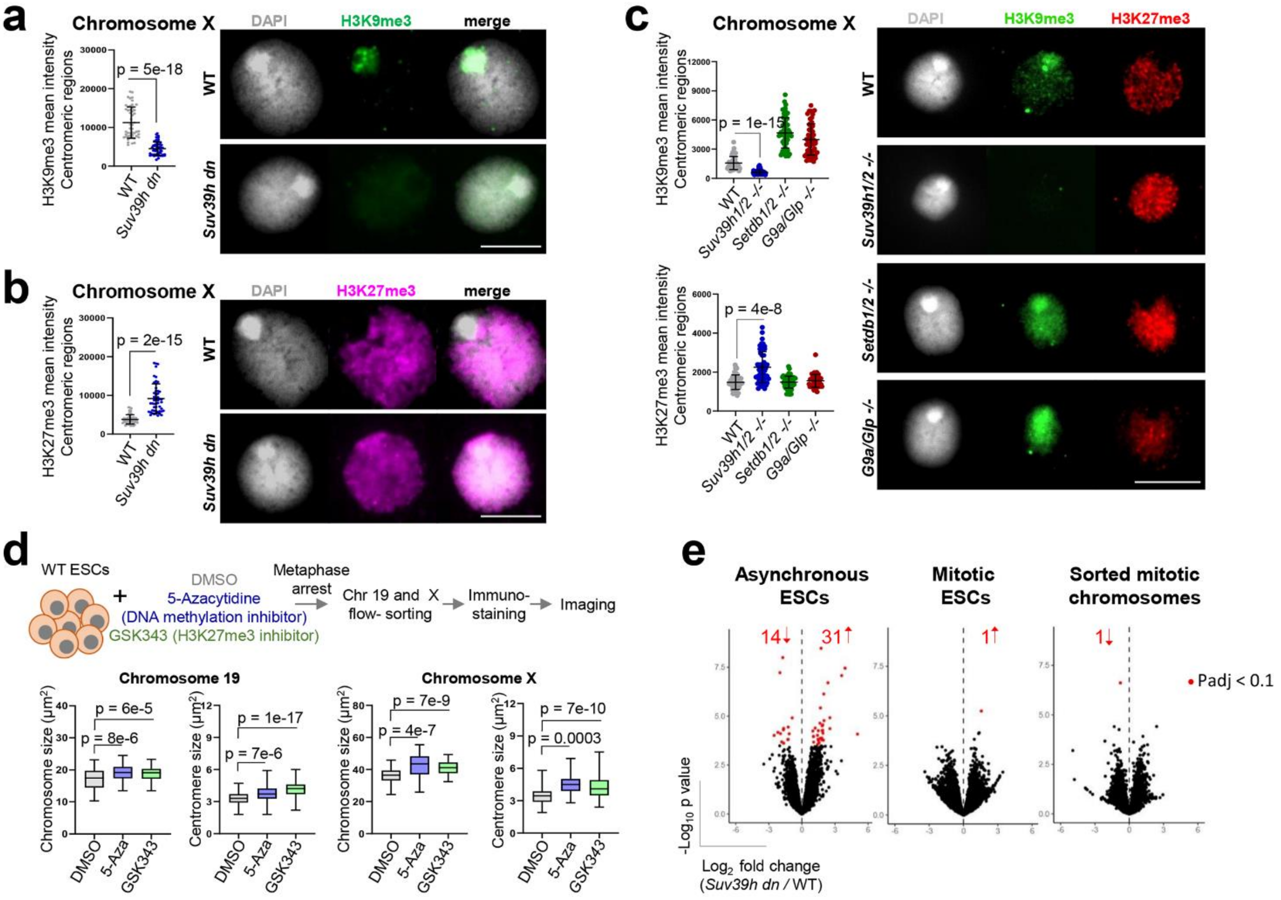


**Figure S2:** (a,b) Representative images (right panel) of immunofluorescence labelling of histone H3K9me3 (a) (green) or histone H3K27me3 (b) (pink) on mouse chromosome X isolated from WT or *Suv39h dn* ESCs where DAPI counterstain is shown in light grey. Scale bars = 5 µm. Plots (left of the images) show H3K9me3 (a) or H3K27me3 (b) mean intensities measured at centromeric regions. Mean ± SD are shown, n = minimum 50 chromosomes over three independent experiments. P-values of statistically significant changes, measured by unpaired two tailed Student's t-tests, are indicated. (c) Representative images (right panel) of histone H3K9me3 (green) and H3K27me3 (red) co-immunolabelling on mouse chromosome X isolated from WT, *Suv39h1/2 -/-* (B1), *Setdb1/2 -/-* (A4+OHT), or *G9a/Glp -/-* (H7) MEFs, where DAPI counterstain is shown in light grey. Scale bars = 5 µm. H3K9me3 mean intensities (upper plot) was measured at centromeric regions for each MEF line, mean ± SD are shown, n = minimum 50 chromosomes over three independent experiments. P-values of statistically significant decreases, measured by unpaired two tailed Student's t-tests, are indicated. H3K27me3 mean intensities (lower plot) was measured at centromeric regions for each MEF line, mean ± SD are shown, n = minimum 50 chromosomes over three independent experiments. P-values of statistically significant increases, measured by unpaired two tailed Student's t-tests, are indicated. (d) Experimental strategy (top panel) used to measure mitotic chromosome size of WT ESCs after treatment with DNA methylation or PRC2 inhibitors (5-Aza or GSK343 respectively). Chromosome and centromere sizes were calculated for each condition. Box plots show area measurements of individual chromosomes and centromeres for each condition. Minimum, lower quartile, median, upper quartile and maximum values are indicated. n = 100 chromosomes over three independent experiments. P-values of statistically significant changes, measured by unpaired two tailed Student's t-tests, are indicated. (e) Volcano plots showing differential accessibility analysis of ATAC-seq peaks for *Suv39h dn* vs WT ESCs.

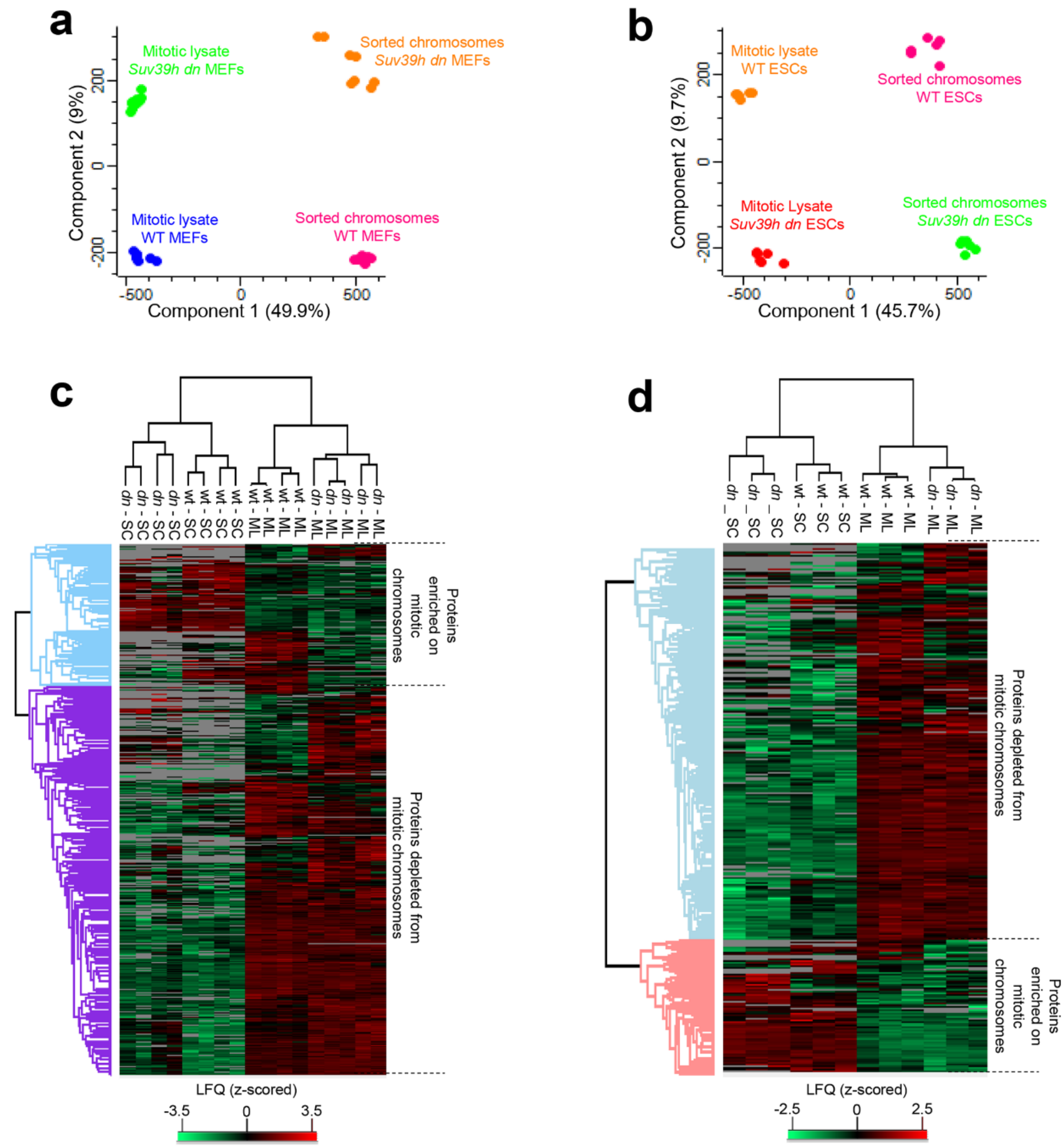


**Figure S3:** (a,b) Principal component analysis (PCA) of proteomic datasets. (c,d) Heatmap and hierarchical clustering analysis (HCA) of significantly changed protein hits (two-sided student's t-test, FDR 0.05) for MEF (c) and ESC (d) samples. Colour scale provided displays z-scored label-free quantification (LFQ) intensities; grey represents missing values (ie not detected in that sample), *dn* = *Suv39h dn*, SC = Sorted chromosomes, ML = Mitotic lysate pellet.

**a**

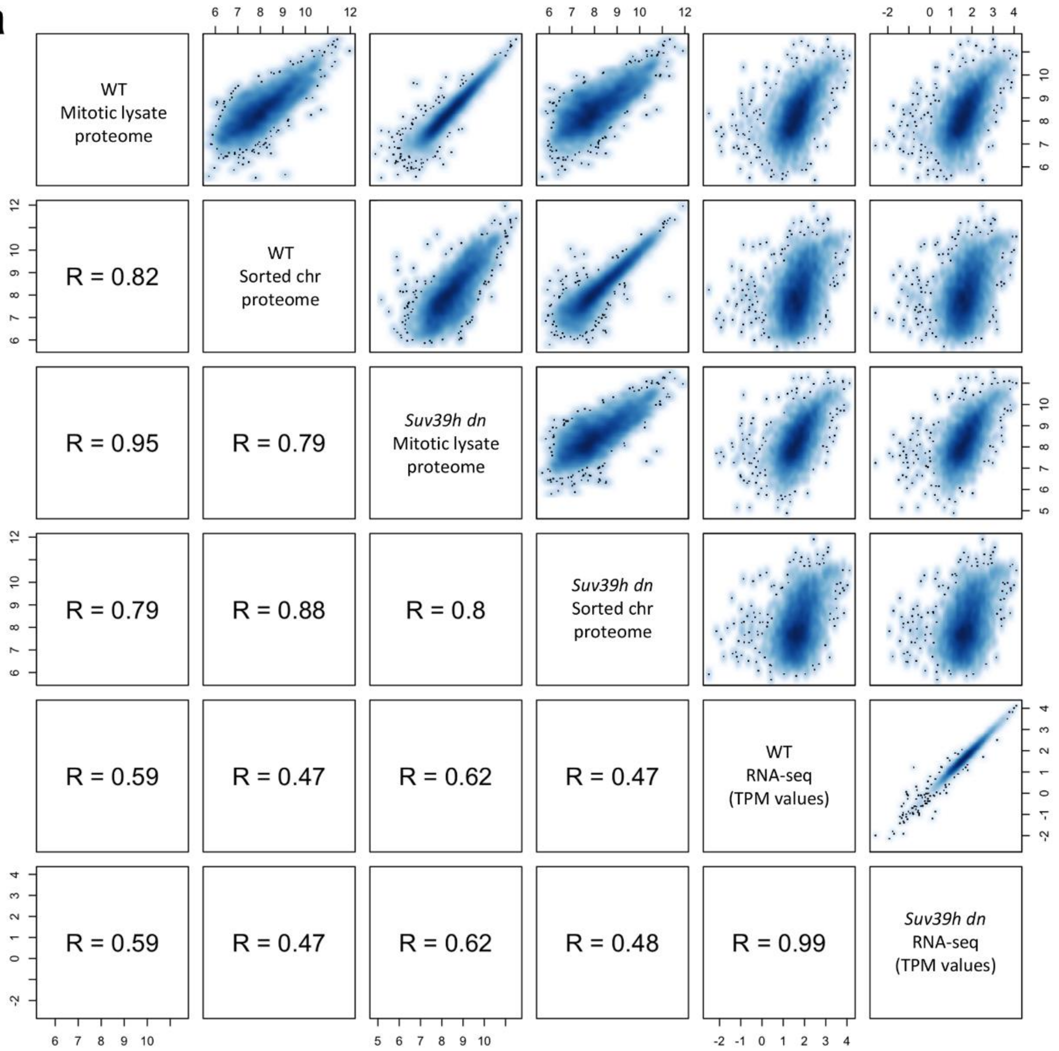
WT
Mitotic lysate
proteome
R = 0.82
WT
Sorted chr
proteome
R = 0.95
R = 0.79
Suv39h dn
Mitotic lysate
proteome
R = 0.79
R = 0.88
R = 0.8
Suv39h dn
Sorted chr
proteome
R = 0.59
R = 0.47
R = 0.62
R = 0.47
WT
RNA-seq
(TPM values)
R = 0.59
R = 0.47
R = 0.62
R = 0.48
R = 0.99
Suv39h dn
RNA-seq
(TPM values)

**b**

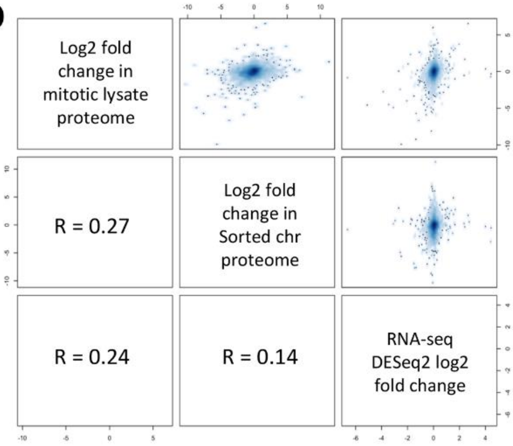
Log2 fold
change in
mitotic lysate
proteome
R = 0.27
Log2 fold
change in
Sorted chr
proteome
R = 0.24
R = 0.14
RNA-seq
DESeq2 log2
fold change

**c**

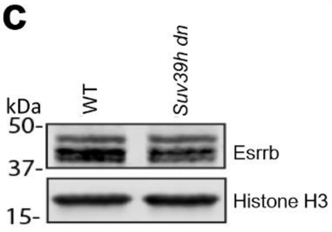
WT
Suv39h dn
kDa
50-
37-
15-
Esrrb
Histone H3

**d**

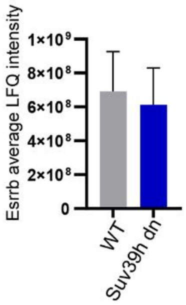
Esrrb average LFQ intensity
1×10⁹
8×10⁸
6×10⁸
4×10⁸
2×10⁸
0
WT
Suv39h dn

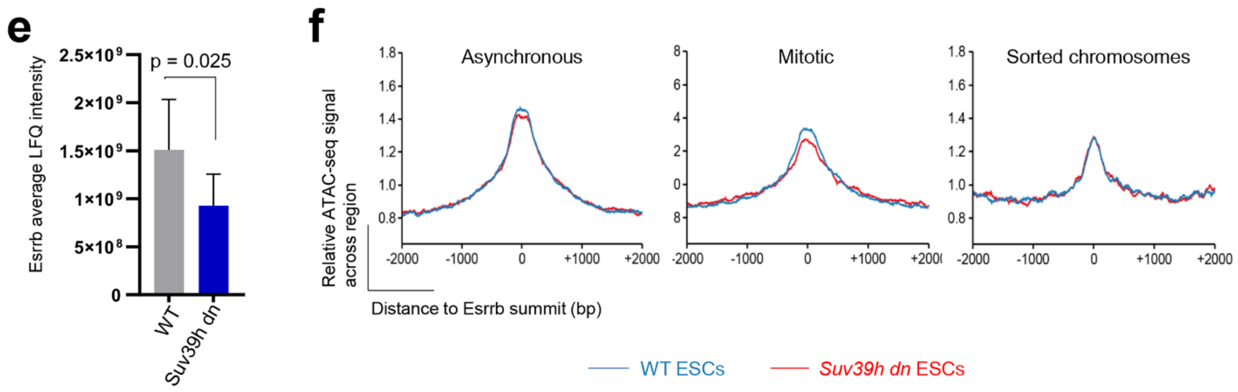


**Figure S4:** (a) Pairwise correlations (R values from Spearman correlation) between proteomics LFQ values (this study) and polyA RNAseq TPM (Transcripts per million) values in ESCs (dataset from[1]). In total, 5664 genes were mapped between proteomics and transcriptomics experiments based on gene symbols. (b) Pairwise correlations (R values from Spearman correlation) between log2 fold changes in proteomics LFQ values (this study) and DESeq2 log2 fold changes in polyA RNAseq values (dataset from[1]) comparing *Suv39h dn* vs WT ESCs. (c) Western blot of Esrrb in WT and *Suv39h dn* asynchronous ESCs. Histone H3 was used as a loading control for the western blot. (d,e) Average LFQ intensities of Esrrb in WT and *Suv39h dn* mitotic lysates (d) and sorted chromosomes (e). Mean + SD is shown, n = 3 independent experiments each measured in duplicate. P-value of statistically significant change, measured by unpaired two tailed Student's t-tests, is indicated. (f) Trend of ATAC-seq accessibility around Esrrb bookmarked binding sites in WT and *Suv39h dn* asynchronous ESCs (left), mitotic ESCs (middle) and sorted chromosomes (right). Esrrb peak locations and bookmarking status are taken from[2].

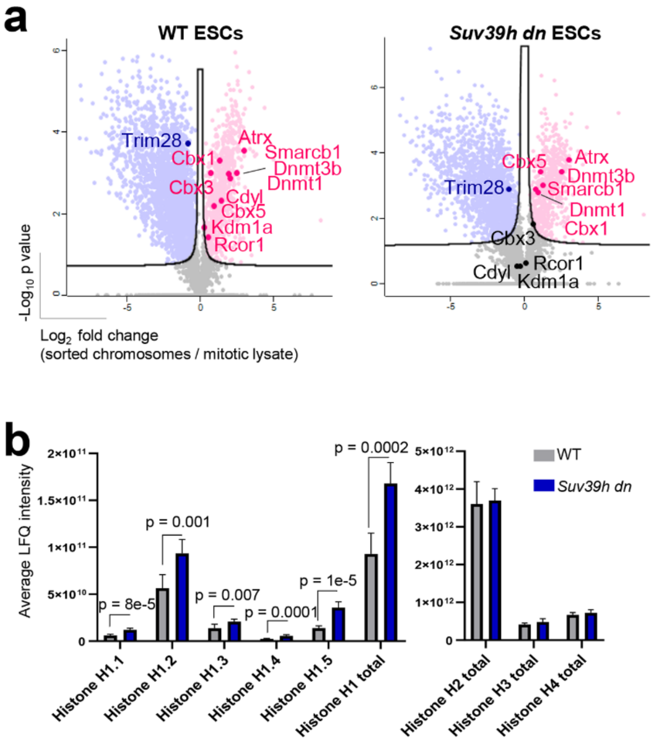


**Figure S5**: (a) Volcano plots as in Figure 3d, highlighting H3K9me3-associated factors that are enriched (red), depleted (blue) or not significantly enriched (black) on WT (left) or *Suv39h dn* (right) ESC mitotic chromosomes versus mitotic lysates. (b) Average LFQ intensity of different histone H1 variants and total histone H1, H2, H3, and H4 in the sorted chromosome samples of WT (grey) and *Suv39h dn* (blue). Mean + SD is shown, n = 3 independent experiments each measured in duplicate. P-values of statistically significant changes, measured by unpaired two tailed Student's t-tests, are indicated.

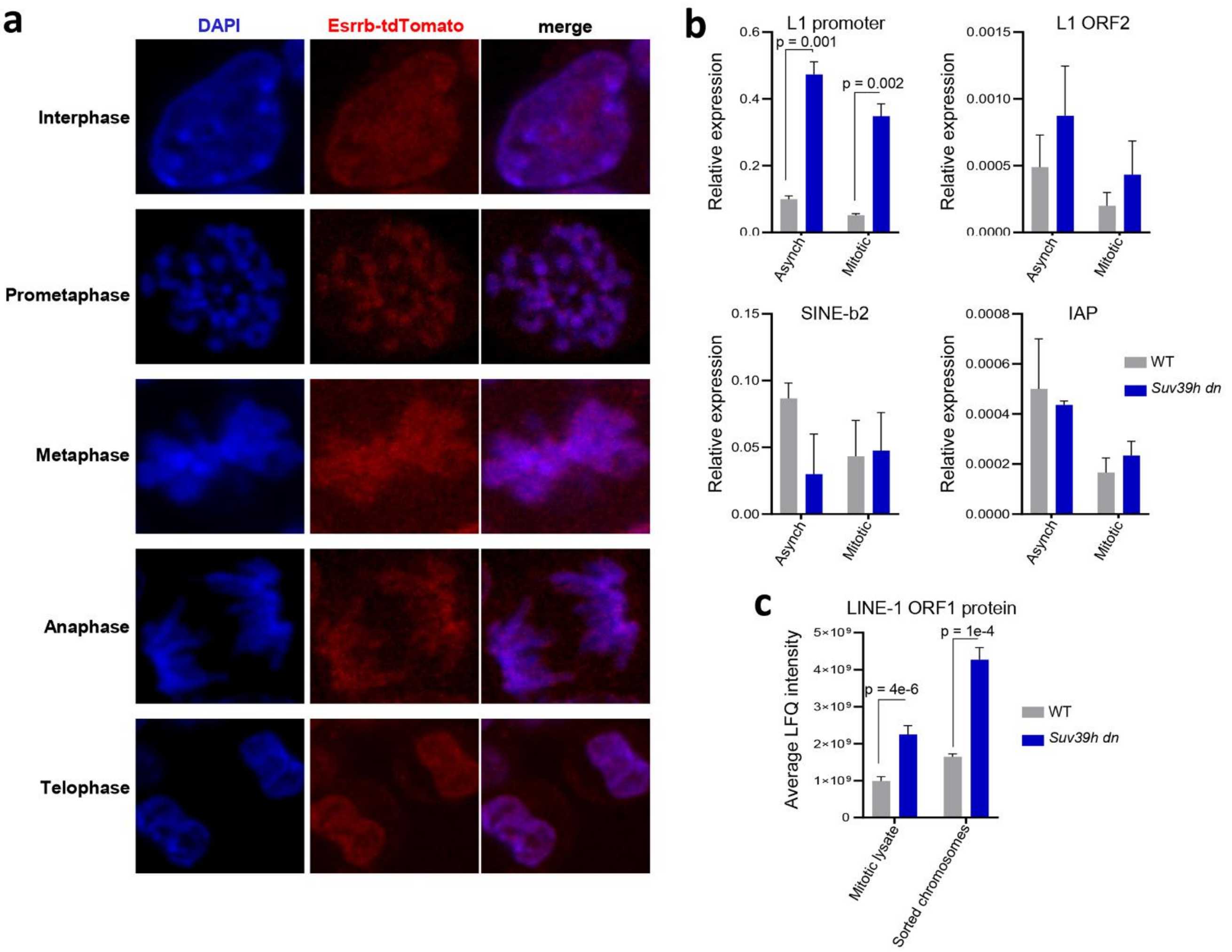


**Figure S6:** (a) Representative images of Esrrb-tdTomato ESCs (DSG+PFA double fixed) in different phases of the cell cycle. DAPI counterstain in blue. Scale bars = 5 µm. (b) qRT-PCR expression analysis of LINE-1, SINE-b2 and IAP repeat elements in WT and *Suv39h dn* asynchronous and mitotic ESCs. Mean + SD is shown, n = 3 independent experiments, P-values of statistically significant changes, measured by unpaired two tailed Student's t-tests, are indicated. (c) Average LFQ intensity of L1ORF1 in mitotic lysates and sorted chromosome samples of WT and *Suv39h dn* ESCs. Mean + SD is shown, n = 3 independent experiments each measured in duplicate. P-values of statistically significant changes, measured by unpaired two tailed Student's t-tests, are indicated.

| ATAC-seq library | Primer name | Primer sequence | Index sequence |
|---|---|---|---|
| All | Ad1 | AATGATACGGCGACCACCGAGATCTACACTCGTCGGCAGCGTCAGATGTG | - |
| WT_Asynchr_1 | Ad2.1 | CAAGCAGAAGACGGCATACGAGATTCGCCTTAGTCTCGTGGGCTCGGAGATGT | TAAGGCGA |
| WT_Asynchr_2 | Ad2.3 | CAAGCAGAAGACGGCATACGAGATTTCTGCCTGTCTCGTGGGCTCGGAGATGT | AGGCAGAA |
| Suv39h dn_Asynchr_1 | Ad2.2 | CAAGCAGAAGACGGCATACGAGATCTAGTACGGTCTCGTGGGCTCGGAGATGT | CGTACTAG |
| Suv39h dn_Asynchr_2 | Ad2.5 | CAAGCAGAAGACGGCATACGAGATAGGAGTCCGTCTCGTGGGCTCGGAGATGT | GGACTCCT |
| WT_Mitotic_1 | Ad2.4 | CAAGCAGAAGACGGCATACGAGATGCTCAGGAGTCTCGTGGGCTCGGAGATGT | TCCTGAGC |
| WT_Mitotic_2 | Ad2.6 | CAAGCAGAAGACGGCATACGAGATCATGCCTAGTCTCGTGGGCTCGGAGATGT | TAGGCATG |
| Suv39h dn_Mitotic_1 | Ad2.7 | CAAGCAGAAGACGGCATACGAGATGTAGAGAGGTCTCGTGGGCTCGGAGATGT | CTCTCTAC |
| Suv39h dn_Mitotic_2 | Ad2.14 | CAAGCAGAAGACGGCATACGAGATACAGTGGTGTCTCGTGGGCTCGGAGATGT | ACCACTGT |
| WT_SC_1 | Ad2.8 | CAAGCAGAAGACGGCATACGAGATCCTCTCTGGTCTCGTGGGCTCGGAGATGT | CAGAGAGG |
| WT_SC_2 | Ad2.19 | CAAGCAGAAGACGGCATACGAGATCCCAACCTGTCTCGTGGGCTCGGAGATGT | AGGTTGGG |
| Suv39h dn_SC_1 | Ad2.9 | CAAGCAGAAGACGGCATACGAGATAGCGTAGCGTCTCGTGGGCTCGGAGATGT | GCTACGCT |
| Suv39h dn_SC_2 | Ad2.20 | CAAGCAGAAGACGGCATACGAGATCACCACACGTCTCGTGGGCTCGGAGATGT | GTGTGGTG |

**Supplemental table 1:** Primers for ATAC-seq library amplification and indexing. Sequences were obtained from[3] and ordered from Sigma-Aldrich (HPLC purified). SC, sorted chromosomes.

| name | F | R | Reference |
|---|---|---|---|
| L1 promoter | ACTGCGGTACATAGGGAAGC | TGTGATCCACTCACCAGAGG | Bulut-Karslioglu et al., (2014)[1] |
| L1 ORF2 | ACCTGGACGAAATGGACAAA | CATCTGGTCCTGGGCTTTT | Bulut-Karslioglu et al., (2014)[1] |
| IAP LTR | AGGGTGGTTCTCTACTCCAT | GAACACCACAGACCAGAATC | Ryu et al., (2011)[4] |
| SINE B2 | GGCTGGTGAGATGGCTCAGT | TACACTGTAGCTGTCTTCAGACA | Allen et al., (2004)[5] |
| β-Actin | CATCCGTAAAGACCTCTATGCCAAC | ATGGAGCCACCGATCCACA | |
| Gapdh | GCGAGACCCCACTAACATCA | CACACCCATCACAAACATGG | |

**Supplemental table 2:** Primers used for quantitative RT-PCR.
